# Unconventional Initiation of PINK1/Parkin Mitophagy by Optineurin

**DOI:** 10.1101/2022.08.14.503930

**Authors:** Thanh Ngoc Nguyen, Justyna Sawa-Makarska, Grace Khuu, Wai Kit Lam, Elias Adriaenssens, Dorotea Fracchiolla, Stephen Shoebridge, Benjamin Scott Padman, Marvin Skulsuppaisarn, Runa S.J. Lindblom, Sascha Martens, Michael Lazarou

## Abstract

Cargo sequestration is a fundamental step of selective autophagy in which cells generate a double membrane structure termed an autophagosome on the surface of cargoes. NDP52, TAX1BP1 and p62 bind FIP200 which recruits the ULK1/2 complex to initiate autophagosome formation on cargoes. How OPTN initiates autophagosome formation during selective autophagy remains unknown despite its importance in neurodegeneration. Here, we uncover an unconventional path of PINK1/Parkin mitophagy initiation by OPTN that does not begin with FIP200 binding nor require the ULK1/2 kinases. Using gene-edited cell lines and *in vitro* reconstitutions, we show that OPTN utilizes the kinase TBK1 which binds directly to the class III phosphatidylinositol 3-kinase complex I to initiate mitophagy. During NDP52 mitophagy initiation, TBK1 is functionally redundant with ULK1/2, classifying TBK1’s role as a selective autophagy initiating kinase. Overall, this work reveals that OPTN mitophagy initiation is mechanistically distinct and highlights the mechanistic plasticity of selective autophagy pathways.

## INTRODUCTION

Macroautophagy (hereafter autophagy) is a highly conserved catabolic process which degrades and recycles intracellular components. Autophagy is cytoprotective and plays essential roles in maintaining cellular homeostasis through the removal of harmful materials such as protein aggregates, dysfunctional organelles, and invading pathogens (Dikic and Elazar, 2018; Melia et al., 2020). Cargoes destined for autophagic degradation are sequestered by double-membrane vesicles termed autophagosomes before being delivered to lysosomes for degradation. Autophagosomes are generated through the formation of a double membrane precursor termed the phagophore that arises on a platform of endoplasmic reticulum termed the omegasome (Axe et al., 2008; Hayashi-Nishino et al., 2009; Uemura et al., 2014).

A general hierarchy of autophagy machinery recruitment during autophagosome formation has been established and typically begins with concurrent recruitment of ATG9A vesicles and the ULK1/2 complex, the latter consisting of the kinase ULK1 or its homologue ULK2, FIP200, ATG13 and ATG101 (Dikic and Elazar, 2018; Itakura and Mizushima, 2010; Kishi-Itakura et al., 2014; Melia *et al*., 2020; Mizushima et al., 2011; Yang and Klionsky, 2010). The ULK1/2 complex then promotes the recruitment and activation of the class III phosphatidylinositol 3-kinase complex I (hereafter PI3K complex). The PI3K complex consists of ATG14, the lipid kinase VPS34, VPS15, and Beclin, which together function to generate the lipid phosphatidylinositol 3-phosphate (PI(3)P) on phagophore membranes. This enables the recruitment of PI(3)P effector proteins including the WIPIs (WD-repeat protein interacting with phosphoinositides) that promote phagophore expansion. WIPI2 plays a role in recruiting the ATG8 conjugation machinery through binding to the ATG16L1 subunit (Dooley et al., 2014), resulting in the attachment of ubiquitin-like ATG8 family proteins onto phosphatidylethanolamine (PE) on autophagosomal membranes (Ichimura et al., 2000; Mizushima et al., 1998). The similarity of ATG8 conjugation to ubiquitination has led to the process being termed atg8ylation, which describes the conjugation of ATG8s to lipid or protein (Agrotis et al., 2019; Deretic and Lazarou, 2022; Kumar et al., 2021; Nguyen et al., 2021). The ATG8 family consists of six members divided equally across the LC3 and GABARAP subfamilies. Attachment of ATG8s to PE may function to expand autophagosomal membranes, at least in part, via promoting membrane tethering and hemifusion (Nakatogawa et al., 2007) and by acting as a protein platform to recruit autophagy factors (Lamark and Johansen, 2021; Martens and Fracchiolla, 2020). WIPI1 and WIPI4 can function to promote membrane expansion through their interactions with ATG2 proteins that transfer phospholipids from the ER to the growing phagophore (Chowdhury et al., 2018; Kotani et al., 2018; Maeda et al., 2019; Osawa et al., 2019; Valverde et al., 2019; Zheng et al., 2017). ATG9A facilitates ATG2-mediated lipid transfer by enabling ER-phagophore contacts and acting as a lipid scramblase by re-arranging lipids from the cytoplasmic leaflet to the luminal leaflet of the growing phagophore (Ghanbarpour et al., 2021; Gomez-Sanchez et al., 2018; Maeda et al., 2020; Matoba et al., 2020). ATG9A vesicles, which are trafficked to autophagosome formation sites by ARFIP2, the immune disease protein LRBA, and the ATG4 family (Judith et al., 2019; Nguyen *et al*., 2021), might also provide the membrane seed for initial phagophore formation (Judith *et al*., 2019; Karanasios et al., 2013; Mari et al., 2010; Sawa-Makarska et al., 2020).

Selective targeting of autophagic cargoes, including the ER, invading pathogens, and mitochondria, necessitates recruitment of autophagy machineries to the surface of cargo to enable selective capture by autophagosomes. Selective recruitment of autophagy machineries is conducted by receptor proteins resident on the surface of cargoes or via adaptor proteins that recognize ‘eat me’ signals on the surface of cargo in the form of ubiquitin chains (Gatica et al., 2018; Goodall et al., 2022; Mizushima, 2020). Several autophagy adaptors have been identified and amongst the most highly studied are p62, NBR1, NDP52, TAX1BP1, and OPTN (Lamark and Johansen, 2021). These autophagy adaptors appear to use a common mechanism of engaging the autophagy machinery by recruiting the ULK1/2 complex through binding to the FIP200 subunit to initiate the cascade of autophagosome formation (Ravenhill et al., 2019; Turco et al., 2021; Turco et al., 2019; Vargas et al., 2019). In addition to the ULK1 and ULK2 kinases, TBK1 is another kinase that has been linked to selective autophagy pathways (Heo et al., 2015; Heo et al., 2018; Lazarou et al., 2015; Moore and Holzbaur, 2016; Richter et al., 2016; Vargas *et al*., 2019), although its precise role during selective autophagy driven by each individual autophagy adaptor remains unclear.

Defects in selective autophagy have been linked to disease in humans, including neurodegeneration in Parkinson’s disease which has been associated with defects in PINK1/Parkin mitophagy (Dikic and Elazar, 2018; Mizushima and Levine, 2020). The kinase PINK1 and the ubiquitin ligase Parkin are mutated in familial Parkinson’s disease (Kitada et al., 1998; Valente et al., 2004) and together function to drive mitophagy of damaged and dysfunctional mitochondria (Kane et al., 2014; Koyano et al., 2014; Narendra et al., 2008; Narendra et al., 2010; Ordureau et al., 2014; Wauer et al., 2015). PINK1 and Parkin generate ubiquitin chains on the mitochondrial outer membrane that recruit the primary mitophagy adaptor proteins, OPTN and NDP52 (Heo *et al*., 2015; Lazarou *et al*., 2015; Wong and Holzbaur, 2014) to initiate autophagosome formation on damaged mitochondria (Lazarou *et al*., 2015; Vargas *et al*., 2019). OPTN and NDP52 are also recruited downstream of autophagosome initiation by ATG8s via an LC3 interacting region (LIR) motif to amplify mitophagy (Padman et al., 2019). To initiate PINK1/Parkin mitophagy, NDP52 directly binds to the FIP200 subunit of the ULK1/2 complex to trigger autophagosome biogenesis (Ravenhill *et al*., 2019; Vargas *et al*., 2019). TBK1 functions to phosphorylate NDP52 and OPTN to increase their affinity for ubiquitin chains and ATG8s (Heo *et al*., 2015; Richter *et al*., 2016; Wild et al., 2011), and also to promote NDP52-FIP200 interactions to drive efficient mitophagy (Vargas *et al*., 2019). In addition, TBK1 coordinates with OPTN for efficient sequestration of damaged mitochondria (Moore and Holzbaur, 2016; Wong and Holzbaur, 2014). However, the mechanism behind how OPTN initiates autophagosome formation during PINK1/Parkin mitophagy remains unclear. A recent *in vitro* study reported the structure of FIP200’s Claw domain bound to the TBK1-phosphorylated LIR domain of OPTN (Zhou et al., 2021), indicating that OPTN may potentially use the same mechanism as NDP52 to initiate PINK1/Parkin mitophagy. Another study reported that an OPTN-ATG9A interaction, that is independent of TBK1, is crucial for PINK1/Parkin mitophagy (Yamano et al., 2020). However, it is unclear if this interaction regulates autophagosome initiation or other downstream steps since ATG9A seems to be involved in multiple stages during autophagosome biogenesis (Gomez-Sanchez et al., 2021). Nevertheless, for autophagy adaptors, the conventional path of selective autophagy initiation identified to date involves direct interaction with FIP200, as has been reported for NDP52 during mitophagy and xenophagy (Ravenhill *et al*., 2019; Vargas *et al*., 2019), and for p62 and TAX1BP1 during aggrephagy (Turco *et al*., 2021; Turco *et al*., 2019).

OPTN is mutated in human neurodegenerative diseases including amyotrophic lateral sclerosis and glaucoma (Maruyama et al., 2010; Rezaie et al., 2002), and unlike NDP52, OPTN is highly expressed in brain tissue (Lazarou *et al*., 2015), making it a high priority cargo adaptor to understand. In this study, we uncover OPTN’s mechanism for initiating selective autophagy. Our work shows that OPTN follows an unconventional path by utilizing TBK1 to interact with the PI3K complex upstream of the ULK1/2 complex. The ULK1/2 kinase subunits were found to be dispensable for OPTN mediated mitophagy, but can contribute to NDP52 mediated mitophagy in a functionally redundant manner with TBK1, revealing TBK1 as a ULK1/2-like kinase in mitophagy. These results demonstrate the mechanistic divergence of mitophagy initiation by OPTN from NDP52 and other cargo adaptors, revealing an alternative path of selective autophagy initiation.

## RESULTS

### ULK1 and ULK2 are dispensable for PINK1/Parkin mitophagy

ULK1 has been reported to phosphorylate several autophagy factors that are important for autophagosome formation (Egan et al., 2015; Mercer et al., 2018; Mercer et al., 2021). Given the role of ULK1 and it relative ULK2 as the most upstream kinases of autophagy, we assessed their contribution to PINK1/Parkin mitophagy initiation mediated by either OPTN or NDP52. Knockouts of ULK1 and ULK2 were conducted in the HeLa penta knockout (KO) background that lacks the five autophagy adaptors OPTN, NDP52, TAX1BP1, NBR1 and p62 (Figure 1A), to enable individual assessment of OPTN- or NDP52-mediated PINK1/Parkin mitophagy. Mitophagy was induced by treating cells with oligomycin and antimycin A (OA) and assessed by measuring the degradation of cytochrome c oxidase subunit II (COXII), a mitochondrial DNA-encoded protein located within the inner membrane. As can be seen in Figures 1B and 1C, the loss of ULK1/2 did not prevent COXII degradation in GFP-NDP52 expressing *ULK1/2* DKO/penta KO cells, and even slightly increased the levels of COXII degradation in GFP-OPTN expressing cells. To determine if the efficiency of mitophagy was affected in the absence of ULK1/2, the mtKeima mitophagy assay was employed (Katayama et al., 2011; Lazarou *et al*., 2015; Vargas *et al*., 2019). Consistent with the COXII degradation data in Figures 1B and 1C, there was no mitophagy defect in penta KO cells lacking ULK1/2 undergoing either GFP-OPTN or GFP-NDP52 driven mitophagy (Figures 1D and 1E). Instead, mitophagy rates appeared to be slightly higher for GFP-OPTN mitophagy (Figure 1D), and to a lesser degree the same was true for GFP-NDP52 (Figure 1E). Analysis of mitochondrial recruitment of the early autophagosome marker WIPI2b showed that the loss of ULK1/2 did not decrease the formation of autophagosome precursors (Figures 1F and 1G). Consistent with the mitophagy measurements in Figures 1D and 1E, the loss of ULK1/2 resulted in more robust formation of WIPI2b autophagosome intermediates during GFP-OPTN-dependent mitophagy and to a lesser degree during GFP-NDP52-dependent mitophagy (Figures 1F and 1G). Autophagosome formation and overall autophagosome morphology in the absence of ULK1/2 also appeared normal by electron microscopy (EM) analysis (Figure 1H). These results show that despite the role of ULK1 and ULK2 as the most upstream kinases of autophagy, they unexpectedly are not essential for autophagosome formation and mitophagy mediated by either OPTN or NDP52.

**Figure 1.**
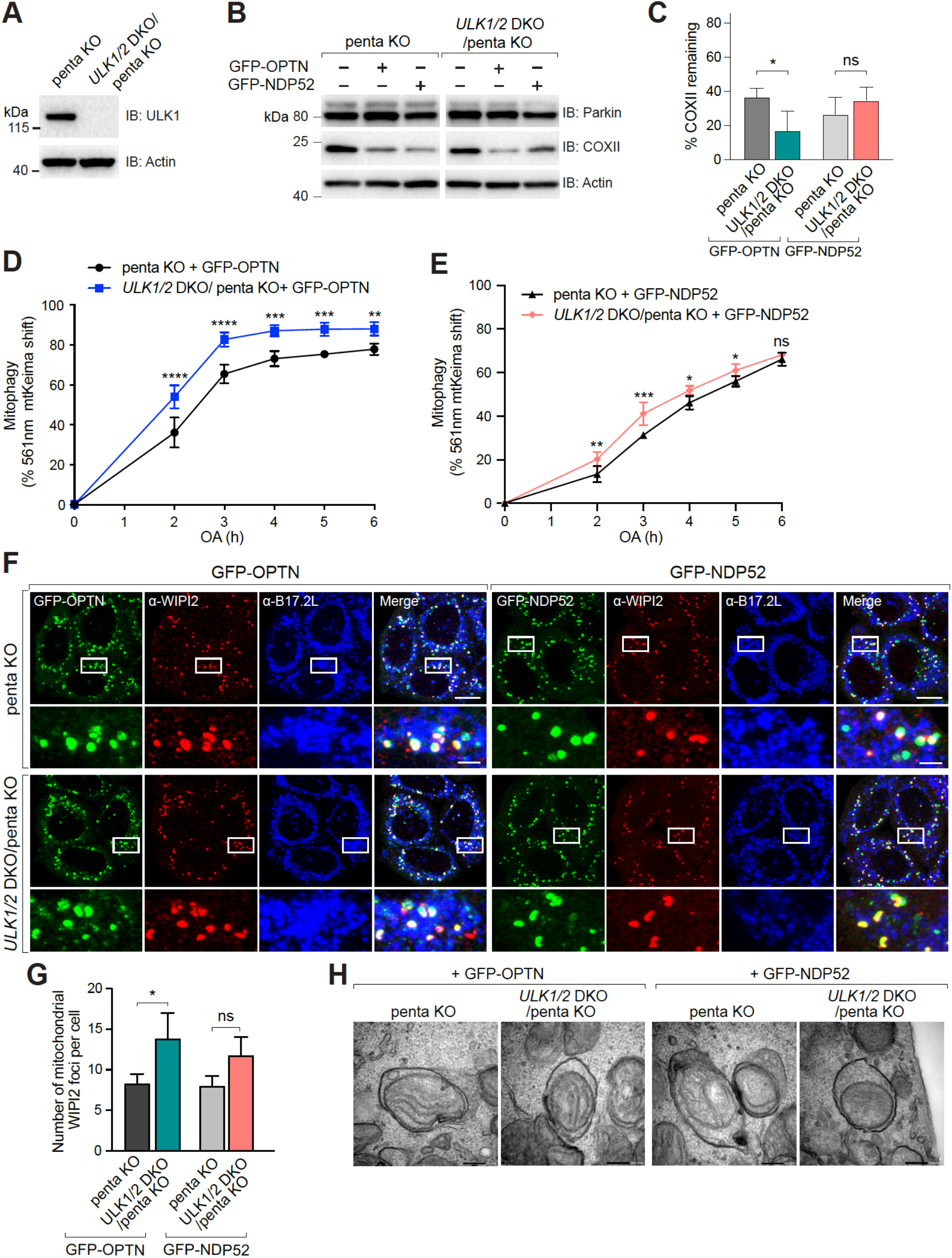
ULK1 and ULK2 are dispensable for Pink1/Parkin mitophagy. (A) *ULK1/2* DKO in penta KO background was confirmed by immunoblotting (IB). kDa: kilodaltons. (B and C) penta KO and *ULK1/2* DKO/penta KO expressing BFP-Parkin and GFP-OPTN or GFP-NDP52 were treated with OA for 18 h and analysed by immunoblotting (B) and CoxII levels were quantified (C). (D and E) penta KO and *ULK1/2* DKO/penta KO cells expressing BFP-Parkin, mtKeima and GFP-OPTN (D) or GFP-NDP52 (E) untreated or treated with OA for indicated times were analysed by FACS. Lysosomal-positive mtKeima was calculated as the percentage of 561 nm mtKeima positive cells. (Representative FACS blots are provided in Figure S1A). (F and G) penta KO and *ULK1/2* DKO/penta KO cells expressing BFP-Parkin and GFP-OPTN or GFP-NDP52 were treated with OA for 1 h and immunostained for WIPI2 and B17.2L (F) and the number of mitochondrial WIPI2 puncta per cell was quantified (G) (Untreated samples shown in Figure S1B). (H) penta KO and *ULK1/2* DKO/penta KO expressing BFP-Parkin and GFP-OPTN or GFP-NDP53 were treated with OA and BafA1 for 3 h and subjected to transmission EM (TEM). Data in (C, D, E and G) are mean ± s.d. from three independent experiments. \**P*<0.05, \*\**P*<0.005, \*\*\**P*<0.001, \*\*\*\**P*<0.0001 (E, G one-way analysis of variance (ANOVA); C, D two-way ANOVA). ns: not significant. Scale bars: (F) overviews, 10 µm; insets, 2 µm; (H) 200 nm.

### The ULK1/2 complex subunits, FIP200 and ATG13, are important for PINK1/Parkin mitophagy

We next asked whether the other subunits of the ULK1/2 kinase complex, including FIP200 and ATG13, are required for PINK1/Parkin mitophagy. Consistent with their essential role in general mammalian autophagy (Hara et al., 2008; Kaizuka and Mizushima, 2016), knockout of FIP200 or ATG13 in penta KO cells (Figures S2A and S2B), blocked PINK1/Parkin mitophagy mediated by either GFP-OPTN or GFP-NDP52, as measured by COXII degradation (Figures 2A and 2B). Next, the recruitment of ATG13 and FIP200 foci to mitochondria was analyzed in penta KO cells lacking FIP200 or ATG13 respectively. In the absence of FIP200, neither GFP-OPTN or GFP-NDP52 could recruit ATG13 foci to mitochondria upon mitophagy induction (Figure 2C). Similarly, in the absence of ATG13, GFP-OPTN could not trigger mitochondrial recruitment of FIP200 foci (Figure 2D). In contrast, GFP-NDP52 was capable of recruiting FIP200 foci to damaged mitochondria in the absence of ATG13 (Figure 2D), which is consistent with the ability of NDP52 to directly bind FIP200 (Ravenhill *et al*., 2019; Vargas *et al*., 2019). These results show that the FIP200 and ATG13 subunits of the ULK1/2 complex are both important for OPTN-and NDP52-dependent PINK1/Parkin mitophagy. Notably, the inability of OPTN to stably recruit FIP200 in the absence of ATG13 indicates that OPTN uses a mechanistically distinct form of mitophagy initiation from NDP52.

**Figure 2.**
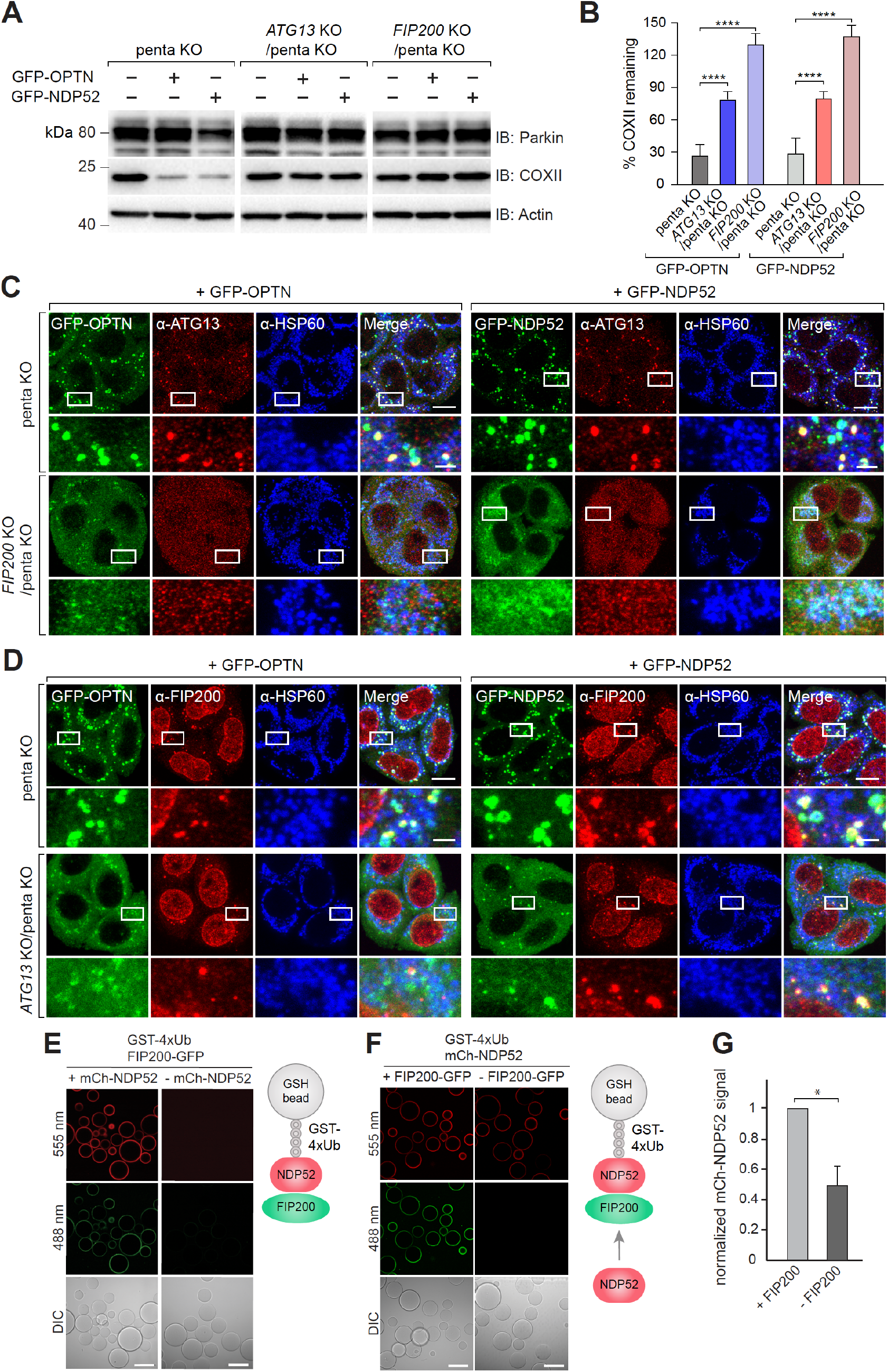
FIP200 and ATG13 are essential for OPTN- and NDP52-mediated mitophagy. (A and B) penta KO, *ATG13* KO/penta KO and *FIP200* KO/penta KO expressing BFP-Parkin and GFP-OPTN or GFP-NDP52 were treated with OA for 18 h and analysed by immunoblotting (A) and CoxII levels were quantified (B). (C) Representative images of penta KO, and *FIP200* KO/penta KO expressing BFP-Parkin and GFP-OPTN or GFP-NDP52 immunostained for ATG13 and mitochondrial HSP60 after 1 h OA treatment. (Untreated images shown in Figure S2C). (D) Representative images of penta KO, and *ATG13* KO/penta KO expressing BFP-Parkin and GFP-OPTN or GFP-NDP52 immunostained for FIP200 and mitochondrial HSP60 after 1 h OA treatment. (Untreated images shown in Figure S2D). (E) Confocal images showing the recruitment of FIP200-GFP to Glutathione sepharose beads coated with GST-tagged linear Ubiquitin chain (GST-4xUb) and incubated with mCherry-NDP52. The cartoon insert depicts the experimental setup. (H and I) Recruitment of mCherry-NDP52 to Glutathione sepharose beads coated with GST-tagged linear Ubiquitin chain (GST-4xUb) in the presence or absence of FIP200-GFP was analysed by confocal imaging (F) and quantified (G). Data in (B) and (G) are mean ± s.d. from three independent experiments. \*\*\**P*<0.001, \*\*\*\**P*<0.0001 (one-way ANOVA; (D)); \**P* ≤ 0.05 (Student’s t-test; (I)). Scale bars in (C) and (D): overviews, 10 µm; insets, 2 µm. Scale bars in (E-F): 100 µm.

In the analyses of GFP-NDP52 mediated recruitment of ATG13 and FIP200 (Figures 2C and 2D), we noted that GFP-NDP52 failed to form mitochondrial foci in the absence of FIP200. This result led us to hypothesize that NDP52-FIP200 interactions can form a positive feedback loop to allow further recruitment and therefore foci formation of NDP52 on damaged mitochondria. We tested this hypothesis by reconstituting NDP52 recruitment *in vitro* using recombinant proteins and a fluorescence microscopy-based bead assay (Abert and Martens, 2019). Recombinant mCherry (mCh)-NDP52 was robustly recruited to cargo mimetic GST-4xUb beads, and this then allowed the recruitment of FIP200-GFP (Figure 2E), providing further confirmation that NDP52 directly interacts with FIP200 (Ravenhill *et al*., 2019; Shi et al., 2020a; Vargas *et al*., 2019). However, in the absence of FIP200, the levels of mCh-NDP52 recruited to the GST-4xUb beads was significantly reduced (Figures 2F and 2G), consistent with the failure to form mitochondrial foci in cells lacking FIP200 (Figure 2C). The presence of FIP200 therefore enhances NDP52 recruitment to cargo, supporting the existence of a FIP200-dependent recruitment loop for NDP52.

### TBK1 is essential for OPTN-mediated mitophagy, but functions redundantly with ULK1/2 during NDP52-mediated mitophagy

TBK1 has been reported to play a role in PINK1/Parkin mitophagy (Heo *et al*., 2015; Heo *et al*., 2018; Lazarou *et al*., 2015; Moore and Holzbaur, 2016; Richter *et al*., 2016; Vargas *et al*., 2019), but its role in mitophagy mediated individually by OPTN or NDP52 has not been explored. This is an important question because the two cargo receptors are not co-expressed in all cell types including neurons. In addition, given the finding that the ULK1/2 kinases are not essential for OPTN or NDP52 mediated mitophagy (Figure 1), further investigation into TBK1 was warranted. First, we assessed the role of TBK1 kinase activity in recruiting the ULK1/2 complex by analyzing mitochondrial recruitment of the ATG13 subunit. Upon mitophagy induction, inhibition of TBK1 kinase activity using BX795 blocked mitochondrial recruitment of ATG13 in penta KO cells undergoing GFP-OPTN mediated mitophagy, but not in cells utilizing GFP-NDP52 for mitophagy (Figure 3A). This result indicates that TBK1 kinase activity is required for initiation of OPTN but not NDP52 mediated mitophagy. For further validation, and to address whether the kinase activity or the physical presence of TBK1 is necessary, TBK1 knockouts were generated in the penta KO background (Figure 3B), and used to analyze mitophagy via COXII degradation, and ULK1/2 complex recruitment via imaging of ATG13. In the absence of TBK1, GFP-OPTN failed to mediate COXII degradation (Figures 3C and 3D) and ATG13 recruitment to mitochondria (Figures 3E and 3F). In contrast, COXII degradation was unaffected by the loss of TBK1 during GFP-NDP52 dependent mitophagy (Figures 3C and 3D), although mitochondrial recruitment of ATG13 was significantly reduced (Figures 3E and 3F). Taken together, these results demonstrate that TBK1 is essential for the recruitment of the ULK1/2 complex and initiation of OPTN-mediated mitophagy, but is largely dispensable for NDP52 mediated mitophagy, although it appears to regulate the efficiency of ULK1/2 complex recruitment and therefore mitophagy initiation by NDP52, as recently reported (Vargas *et al*., 2019).

**Figure 3.**
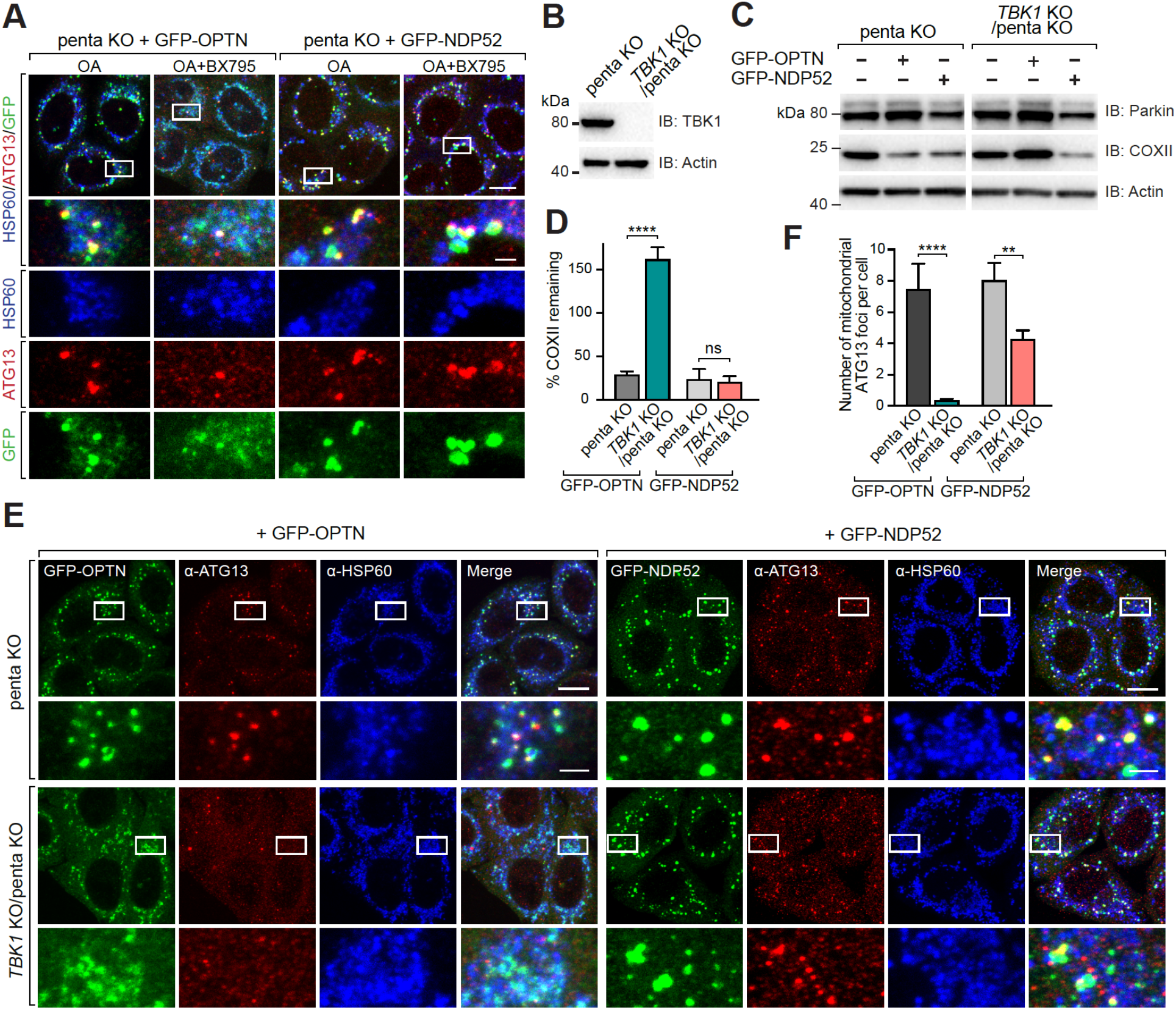
TBK1 is essential for OPTN-mediated mitophagy but not NDP52-mediated mitophagy. (A) penta KO cells expressing GFP-OPTN or GFP-NDP52 were treated with OA or OA and BX795 for 1 h and immunostained for ATG13 and mitochondrial HSP60. (B) *TBK1* KO/penta KO was confirmed by immunoblotting (IB). kDa: kilodaltons. (C and D) penta KO and *TBK1* KO/penta KO expressing BFP-Parkin and GFP-OPTN or GFP-NDP52 were treated with OA for 18 h and analysed by immunoblotting (C) and CoxII levels were quantified (D). (E and F) penta KO and *TBK1* KO/penta KO expressing BFP-Parkin and GFP-OPTN or GFP-NDP52 treated with OA for 1 h and immunostained for ATG13 and mitochondrial HSP60 (E) and the number of mitochondrial ATG13 foci per cell was quantified (F). (Untreated images shown in Figure S3A). Data in (D and F) are mean ± s.d. from three independent experiments. \**P*<0.05, \*\**P*<0.005, \*\*\**P*<0.001, \*\*\*\**P*<0.0001 (one-way ANOVA). ns: not significant. Scale bars: overviews, 10 µm; insets, 2 µm.

Neither of the ULK1/2 or TBK1 kinases were essential for NDP52-mediated mitophagy (Figures 1-3). However, given that some structural similarities between TBK1 and the ULK1/2 complex have been reported (Shi et al., 2020b), we asked whether TBK1 and ULK1/2 might function redundantly during NDP52 mediated mitophagy. Mitophagy rates using the mtKeima assay were analyzed in *ULK1/2* DKO/penta KO cells expressing either GFP-OPTN or GFP-NDP52 in the presence or absence of TBK1 inhibitor BX795. GFP-OPTN mediated mitophagy was completely inhibited upon addition of BX795 irrespective of the presence or absence of ULK1/2 (Figures 4A) supporting the absolute requirement of TBK1 kinase activity for OPTN dependent mitophagy. In contrast, TBK1 kinase inhibition with BX795 was only able to prevent NDP52 dependent mitophagy when ULK1/2 were absent (Figure 4B). This demonstrates that TBK1 kinase activity compensates for the loss of ULK1/2 during NDP52 mediated mitophagy.

**Figure 4.**
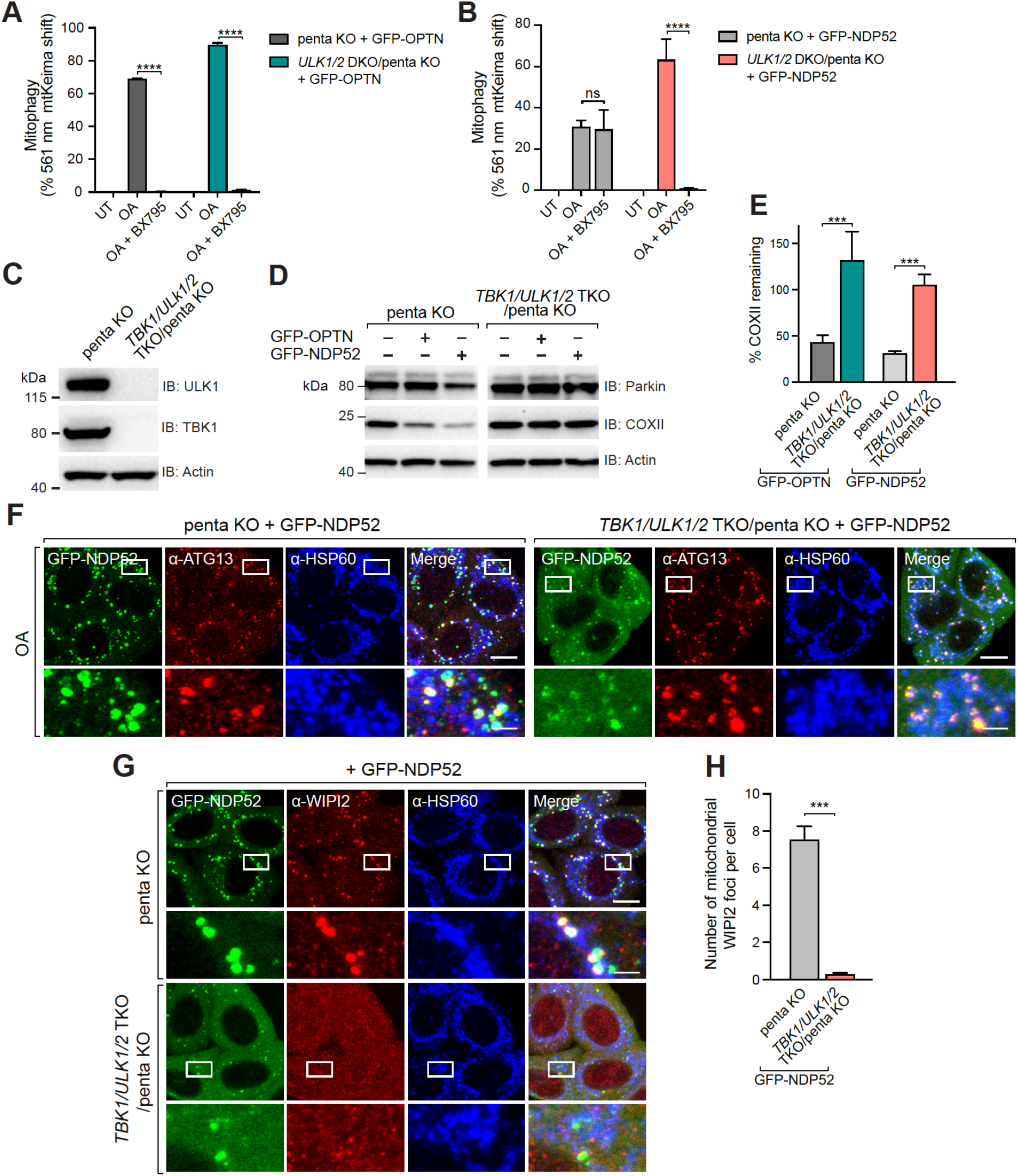
TBK1 and ULK1/2 function redundantly during NDP52-dependent mitophagy. (A and B) penta KO and *ULK1/2* DKO/penta KO cells expressing BFP-Parkin, mtKeima and GFP-OPTN (A) or GFP-NDP52 (B) untreated or treated with OA or OA and BX795 for 4 h were analysed by FACS. Lysosomal-positive mtKeima was calculated as the percentage of 561 nm mtKeima positive cells. (Representative FACS blots are provided in Figure S3B). (C) *TBK1/ULK1/2* TKO/penta KO was confirmed by immunoblotting (IB). kDa: kilodaltons. (D and E) penta KO and *TBK1/ULK1/2* TKO/penta KO expressing BFP-Parkin and GFP-OPTN or GFP-NDP52 were treated with OA for 18 h and analysed by immunoblotting (D) and CoxII levels were quantified (E). (F) penta KO and *TBK1/ULK1/2* TKO/penta KO cells expressing BFP-Parkin and GFP-NDP52 were immunostained for ATG13. (G and H) penta KO and *TBK1/ULK1/2* TKO/penta KO cells expressing BFP-Parkin and GFP-NDP52 were immunostained for WIPI2 and mitochondrial HSP60 (G) and the number of mitochondrial WIPI2 foci per cell was quantified (H). (Representative images of untreated samples shown in Figure S3C and S3D). Data in (A, B, E and H) are mean ± s.d. from three independent experiments. \**P*<0.05, \*\**P*<0.005, \*\*\**P*<0.001, \*\*\*\**P*<0.0001 (A, B two-way analysis of variance (ANOVA); E and H one-way ANOVA). ns: not significant. Scale bars: overviews, 10 µm; insets, 2 µm.

To further address the putative functional redundancy between TBK1 and ULK1/2 during NDP52 mediated mitophagy initiation, and to ensure no off-target effects of BX795 on other kinases, TBK1 was knocked out in the *ULK1/2* DKO/penta KO background (Figure 4C). The combined deletion of TBK1 and ULK1/2 resulted in GFP-NDP52 losing its ability to drive COXII degradation (Figures 4D and 4E), demonstrating that indeed, ULK1/2 and TBK1 are functionally redundant in initiating NDP52 mediated mitophagy. Analysis of ATG13 recruitment to mitochondria showed that consistent with its direct binding to FIP200, GFP-NDP52 was still able to support recruitment of ATG13 in the absence ULK1/2 and TBK1 (Figure 4F), demonstrating that a core complex consisting of FIP200, ATG13, and likely ATG101, remains intact (Jung et al., 2009; Shi *et al*., 2020b). However, the recruitment of WIPI2b, a PI(3)P effector protein, was completely abolished (Figures 4G and 4H), pointing toward a defect in the activation or recruitment of the PI3K complex. Our results therefore indicate that TBK1 is functionally redundant with ULK1/2 in promoting the activation of PI3K complex, which is consistent with the role of ULK1/2 in enhancing activity of the PI3K complex during starvation induced autophagy (Mercer *et al*., 2018; Mercer *et al*., 2021; Wold et al., 2016).

### OPTN initiates mitophagy through an unconventional mechanism that is dependent on TBK1

A recent structural analysis revealed that peptides of OPTN and FIP200 interact *in vitro* (Zhou *et al*., 2021), indicating that OPTN follows the canonical mode of selective autophagy activation of directly recruiting the ULK1/2 complex via FIP200. However, our results show that OPTN is unable to stably recruit FIP200 in the absence of ATG13 (Figure 2D), arguing against FIP200 binding being the primary mitophagy initiation mechanism of OPTN. To directly address whether FIP200 binding, and therefore ULK1/2 complex recruitment, is the most upstream event of OPTN mitophagy initiation, we decided to block the next step of mitophagy involving PI3K complex recruitment. The rationale being that if the ULK1/2 complex is the most upstream event, then OPTN should retain the ability to recruit the ULK1/2 complex irrespective of downstream mitophagy steps. Knockout lines of ATG14, an essential subunit of the PI3K complex involved in autophagy, were generated in the penta KO background (Figure 5A). Analysis of COXII degradation confirmed that ATG14 is essential for PINK1/Parkin mitophagy mediated by either GFP-OPTN or GFP-NDP52 (Figures 5B and 5C). We also confirmed that the loss of ATG14 blocked activation of the PI3KC3 complex as shown by the lack of mitochondrial recruitment of the PI(3)P effector protein WIPI2b (Figure 5F).

**Figure 5.**
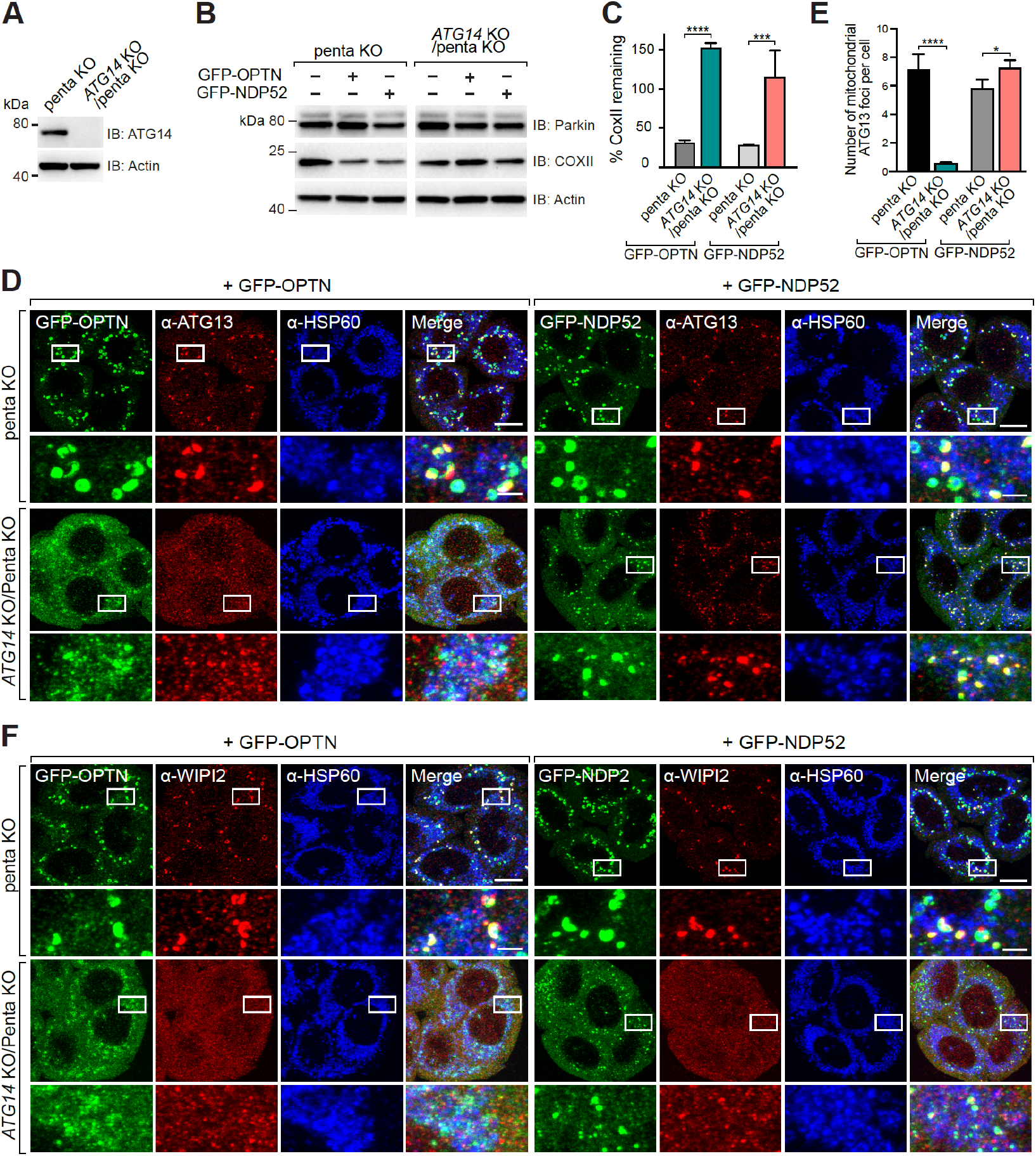
ATG14 is crucial for mitochondrial recruitment of ATG13/FIP200 complex during OPTN-dependent mitophagy. (A) *ATG14* KO in penta KO background was confirmed by immunoblotting (IB). kDa: kilodaltons. (B and C) penta KO and *ATG14* KO/penta KO expressing BFP-Parkin and GFP-OPTN or GFP-NDP52 were treated with OA for 18 h and analysed by immunoblotting (B) and CoxII levels were quantified (C). (D-F) penta KO and *ATG14* KO/penta KO expressing BFP-Parkin and GFP-OPTN or GFP-NDP52 were treated with OA for 1 h and immunostained for ATG13 and mitochondrial HSP60 (D) or WIPI2 and mitochondrial HSP60 (F). The number of mitochondrial ATG13 foci per cell in (D) was also quantified (E). (Representative images of untreated samples shown in Figure S4A and S4C). Data in (C) and (E) are mean ± s.d. from three independent experiments. \**P*<0.05, \*\**P*<0.005, \*\*\**P*<0.001, \*\*\*\**P*<0.0001 (one-way analysis of variance (ANOVA)). ns: not significant. Scale bars: overviews, 10 µm; insets, 2 µm.

Next, the recruitment of the ULK1/2 complex was analyzed by assessing ATG13 and FIP200 foci on mitochondria in the presence or absence of ATG14. During GFP-NDP52 mediated mitophagy, both ATG13 and FIP200 were recruited to damaged mitochondria irrespective of the presence or absence of ATG14 (Figures 5D, 5E and S4B), aligning with NDP52’s ability to directly bind and recruit FIP200 upstream of the PI3K complex (Ravenhill *et al*., 2019; Vargas *et al*., 2019). However, in direct contrast, cells undergoing GFP-OPTN mediated mitophagy failed to recruit ATG13 and FIP200 in cells lacking ATG14 (Figures 5D, 5E and S4B). This result argues for an unconventional mechanism of mitophagy activation by OPTN, in which the PI3K complex is upstream of the ULK1/2 complex. Combined with the finding that TBK1 is essential for OPTN but not NDP52 mediated mitophagy, we conclude that mitophagy initiation by OPTN is mechanistically distinct from NDP52.

### TBK1 directly binds to the PI3K complex

We next investigated whether OPTN can directly engage the PI3K complex by reconstituting mitophagy initiation *in vitro*. Ubiquitin coated cargo mimetic beads were incubated with a TBK1 phospho-mimetic form of OPTN, termed OPTN S2D (S177D, S473D (Chang et al., 2021)), that was used to assess recruitment of the PI3K complex. Phosphomimetic OPTN S2D failed to recruit fluorescently labeled PI3K complex (mCherry-ATG14) to GST4xUb beads (Figure 6A), arguing against a direct binding model for OPTN mitophagy initiation. However, the PI3K complex was recruited upon the addition of GFP-TBK1 (Figure 6A). This led us to assess whether TBK1 can directly bind to the PI3K complex. Indeed, GFP-TBK1 alone tethered to GFP-Trap beads was sufficient to recruit the PI3K complex (Figure 6B). *In vitro* reconstitution of NDP52 mediated mitophagy initiation using ubiquitin coated beads was conducted next. The analysis showed that that NDP52 can directly recruit the ULK1 complex (Figure 6C), and this was sufficient to then recruit the PI3K complex (Figure 6D), indicating that purified ULK1 and PI3K complexes alone are sufficient for the two to directly interact. Another *in vitro* reconstitution in which the PI3K complex was immobilized onto RFP-Trap beads via mCherry-ATG14 confirmed that a direct biochemical interaction exists between the ULK1 and PI3K complexes (Figure S5A).

**Figure 6.**
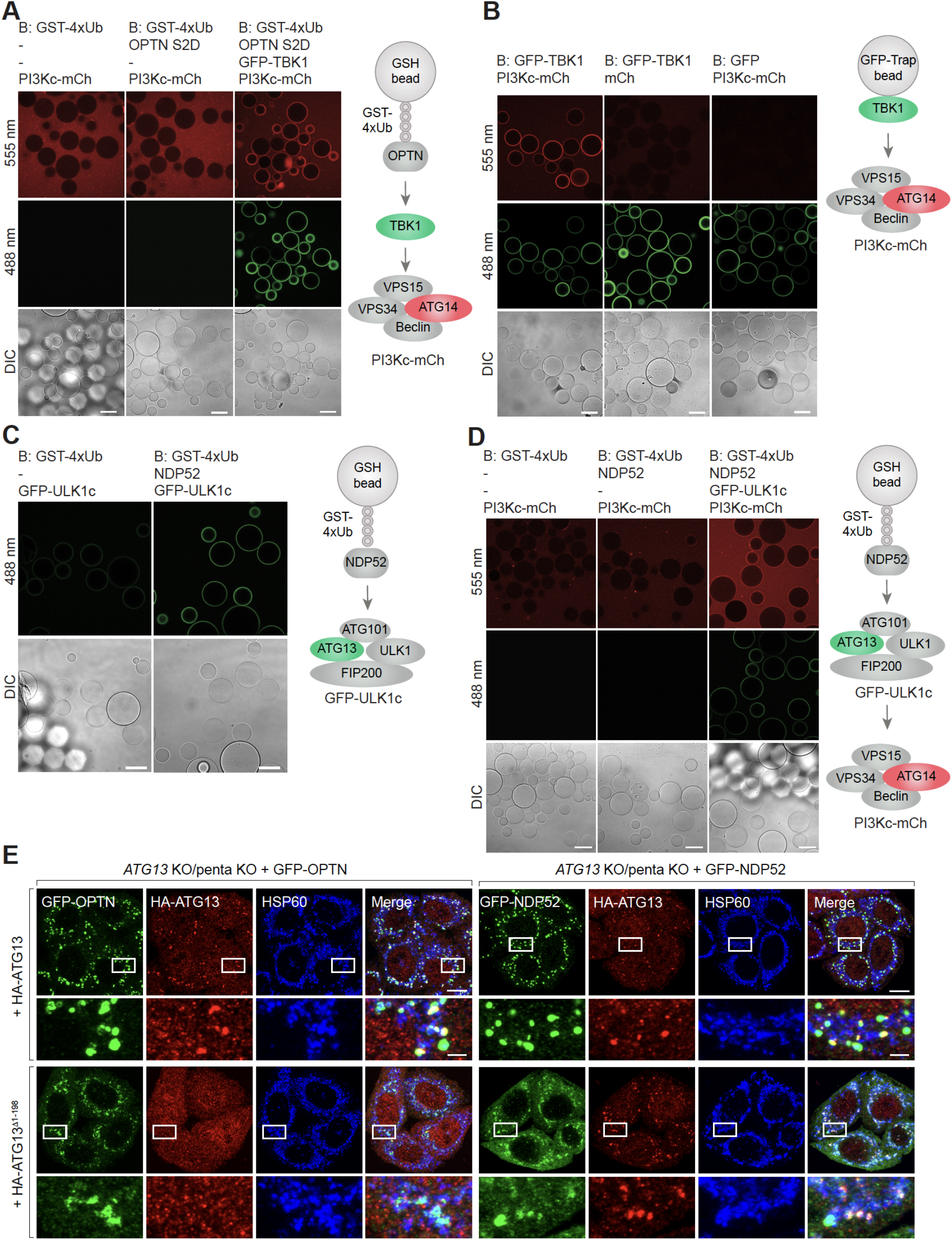
PI3K complex directly interacts with TBK1 to allow mitochondrial recruitment of ATG13 during OPTN-mediated mitophagy. (A) Confocal images showing recruitment of GFP-TBK1 and PI3Kc1-mCh on cargo mimetic beads. Glutathione sepharose (GSH) beads were coated with GST-tagged linear Ubiquitin chain (GST-4xUb) and incubated with indicated protein components and visualized by confocal microscope (B) Microscopy-based bead protein interaction assay with GFP-trap beads using GFP or GFP-TBK1 as baits and mCherry (mCh) or PI3K complex-mCherry (PI3Kc-mCh) as preys. (C) Representative confocal images of GFP-ULK1 complex (GFP-ULK1c) recruitment to NDP52 bound to GST-tagged linear Ubiquitin chain (GST-4xUb). (D) Representative confocal images of PI3K complex-mCherry (PI3Kc-mCh) recruitment to GST-4xUb/NDP52 axis via ULK1c-GFP. The cartoons in (A-D) depict the experimental setups. (E) *ATG13* KO/penta KO cells expressing BFP-Parkin and GFP-OPTN or GFP-NDP52 rescued with HA-ATG13 or HA-ATG13^.1HORMA^ were treated with OA for 1 h and immunostained for HA and mitochondrial HSP60. (Untreated images shown in Figure S5B). Scale bars: (A-D) 100 µm; (D) overviews, 10 µm; insets, 2 µm.

The positioning of the PI3K complex upstream of the ULK1/2 complex during OPTN mediated mitophagy raised the question of how the ULK1/2 complex might be recruited by OPTN. Given that the PI3K and ULK1 complexes directly bind *in vitro* (Figure S5A) and have been reported to interact in cells (Park et al., 2016), it was decided to explore this area further. Using human cells, it has been proposed that the N-terminal HORMA domain within ATG13 interacts with ATG14 (Park *et al*., 2016), while in yeast it plays a role in the recruitment of ATG14 (Jao et al., 2013). We therefore assessed whether the HORMA domain of ATG13 is required for recruitment of the ULK1/2 complex during OPTN mediated mitophagy. As can be seen in Figure 6E, deletion of the HORMA domain within ATG13 (ATG13^Δ1-198^) prevented its recruitment to mitochondria during OPTN, but not NDP52, mediated mitophagy (Figure 6E). The ability of NDP52 to recruit ATG13^Δ1-198^ is explained by NDP52’s direct binding to FIP200 which interacts with the C-terminus of ATG13 (Jung *et al*., 2009; Shi *et al*., 2020b). Collectively, our results demonstrate that OPTN initiates autophagosome formation via direct interaction between TBK1 and the PI3K complex, whereas NDP52 initiates mitophagy via direct interaction with the ULK1/2 complex (Figure S5C), while the HORMA domain of ATG13 plays a linking role between the PI3K and ULK1/2 complexes (Figure S5C). The linking role played by ATG13 explains how OPTN’s unconventional mechanism of mitophagy initiation can proceed by first engaging with the PI3K complex through TBK1.

## DISCUSSION

During PINK1/Parkin mitophagy, OPTN and NDP52 enable the recruitment of early autophagy machineries that initiate the *de novo* formation of autophagosomes on the surface of damaged mitochondria (Lazarou *et al*., 2015; Vargas *et al*., 2019). The mechanism of NDP52-mediated mitophagy initiation was recently revealed and involves NDP52 binding to the C-terminal part of FIP200 via an N-terminal SKICH domain (Vargas *et al*., 2019). The interaction between NDP52 and FIP200 enables recruitment and activation of the ULK1/2 complex through local clustering (Vargas *et al*., 2019). An analogous mechanism was identified during xenophagy for NDP52 (Ravenhill *et al*., 2019), and for TAX1BP1 and p62 during aggrephagy (Turco *et al*., 2021; Turco *et al*., 2019). The discovery that the LIR region of OPTN can also bind to the Claw domain of FIP200 *in vitro* indicated that OPTN also initiates selective autophagy in the same way. However, OPTN has also been reported to bind ATG9A in cells (Yamano *et al*., 2020), and to date it has remained unclear exactly how OPTN initiates selective autophagy. The importance of understanding OPTN’s initiation mechanism is highlighted by the fact that it is mutated in amyotrophic lateral sclerosis (ALS) (Maruyama *et al*., 2010), is involved PINK1/Parkin mitophagy linked to Parkinson’s disease (Kitada *et al*., 1998; Valente *et al*., 2004; Youle, 2019), and is abundantly expressed in brain tissue where there is little to no NDP52 (Lazarou *et al*., 2015).

By delineating the initiation mechanism of OPTN in isolation during PINK1/Parkin mitophagy, we have revealed an unconventional pathway of selective autophagy initiation that is independent of FIP200 binding as the most upstream event. We propose a model in which OPTN’s binding partner TBK1 (Li et al., 2016; Richter *et al*., 2016), recruits/activates the PI3K complex via direct binding, which is linked to ULK1/2 complex via the HORMA domain within ATG13 (Figures 6E and S5C), thereby bringing the ULK1/2 complex to the surface of mitochondria. Upon recruitment of the ULK1/2 complex, OPTN’s interaction with FIP200 (Zhou *et al*., 2021) may become stabilized, and in combination with the recruitment of ATG9A vesicles via direct interaction with OPTN (Yamano *et al*., 2020), the formation of phagophore membranes is triggered (Figure 6F). The presence of ATG9A vesicles at this early stage likely provides lipids for generating the phagophore (Judith *et al*., 2019; Karanasios *et al*., 2013; Mari *et al*., 2010; Sawa-Makarska *et al*., 2020; Yamamoto et al., 2012). Whether the complex interplay of OPTN mediated interactions is mutually exclusive to NDP52 mitophagy initiation remains to be determined. While our data supports that engagement of the PI3K complex occurs first via a direct interaction with TBK1, the recruitment of the PI3K and ULK1/2 complex to autophagosome formation site is likely to be more complex and dynamic. Indeed, previous work has identified oscillatory recruitment of ATG13 and OPTN, as well as positive feedback loops between PI3K and ULK1/2 complex recruitment (Karanasios *et al*., 2013; Zachari et al., 2019). Whether TBK1 mediated phosphorylation of the PI3K and ULK1/2 complexes regulates the dynamics of initiation (Figure S5D), including OPTN binding to FIP200, warrants further investigation.

Given the essential role played by ULK1/2 kinases in starvation autophagy initiation (Chan et al., 2007; Hosokawa et al., 2009; Jung *et al*., 2009), it was reasonable to conclude that they should be essential for PINK1/Parkin mitophagy. Instead, we discovered that ULK1/2 are dispensable for mitophagy initiated by either OPTN or NDP52. More specifically, ULK1/2 are completely dispensable for OPTN mediated mitophagy, but are functionally redundant with TBK1 during NDP52 mediated mitophagy (Figure 4), although mitophagy through TBK1 appears to be slightly faster (Figure 1). A PINK1/Parkin independent form of mitophagy induced by ivermectin treatment also does not require ULK1/2 and is instead driven by TBK1 (Zachari *et al*., 2019), while other autophagy pathways independent of ULK1/2 kinase activity have also been identified (Alers et al., 2011; Hieke et al., 2015). It is possible that TBK1 compensates for the loss of ULK1/2 in the ULK1/2 independent pathways. We propose that TBK1 can function as a selective autophagy initiating kinase, akin to ULK1/2, but TBK1’s role in selective autophagy will depend on which autophagy adaptor is involved. For example, TBK1 can play a dominant role during OPTN mediated selective autophagy through stable direct binding (Li *et al*., 2016; Richter *et al*., 2016). While the ULK1/2 kinases were not essential for PINK1/Parkin mitophagy (Figure 1), the ATG13 and FIP200 subunits were found to play an important role (Figure 2), highlighting the importance of key subunits of the ULK1/2 complex. During NDP52 mediated mitophagy, FIP200 was found to promote the recruitment of NDP52 both in cells and *in vitro* (Figure 2) in an apparent positive feedback loop. The discovery that membrane binding drives allosteric activation of ULK1/2 complex recruitment by FIP200 and NDP52 likely also contributes to the positive feedback loop in cells (Shi *et al*., 2020a).

In yeast, the HORMA domain of Atg13 was shown to be important for Atg14 recruitment to autophagosome formation sites during autophagy (Jao *et al*., 2013). In mammalian systems, the PI3K complex has also been reported interact with the ULK1 complex and this interaction is mediated by direct binding between ATG13 and ATG14 via the HORMA domain in ATG13 (Park *et al*., 2016). OPTN and NDP52 mediated mitophagy is consistent with the HORMA domain of ATG13 playing a role in connecting the PI3K and ULK1/2 complexes (Figure 6) which is crucial for ULK1/2 complex recruitment during OPTN mediated mitophagy. It would be interesting to investigate the role of the HORMA domain within ATG101, an ULK1/2 complex subunit that has been shown to dimerize with ATG13 (Qi et al., 2015; Suzuki et al., 2015). ATG101 could conceivably allow the simultaneous binding of the ULK1/2 complex to ATG9A and the PI3K complex to enhance autophagosome formation (Ren et al., 2022).

Overall, our discovery that OPTN and NDP52 utilise distinct mechanisms to drive mitophagy initiation has broad implications for selective autophagy. It indicates that within the same type of selective autophagy, multiple mechanisms of initiation can be undertaken depending on which autophagy adaptors are present, and that can vary across different cell types and tissues. This finding can become important when it comes to targeting a particular selective autophagy pathway for therapeutic purposes. For example, strategies to enhance PINK1/Parkin mitophagy in neurons that express high levels of OPTN will differ from that in tissues that express little to no OPTN, such as heart and small intestine (Lazarou *et al*., 2015). It would also be important to dissect whether the mechanism of mitophagy initiation driven by OPTN is more suited to the cell biology of a neuron. Digging deeper into the precise mechanisms of selective autophagy initiation mediated by each autophagy adaptor during different forms of selective autophagy is likely to yield therapeutic strategies to target damaged mitochondria, invading pathogens, and aggregating proteins in human disease.

## Supplementary Data

**Figure S1.**
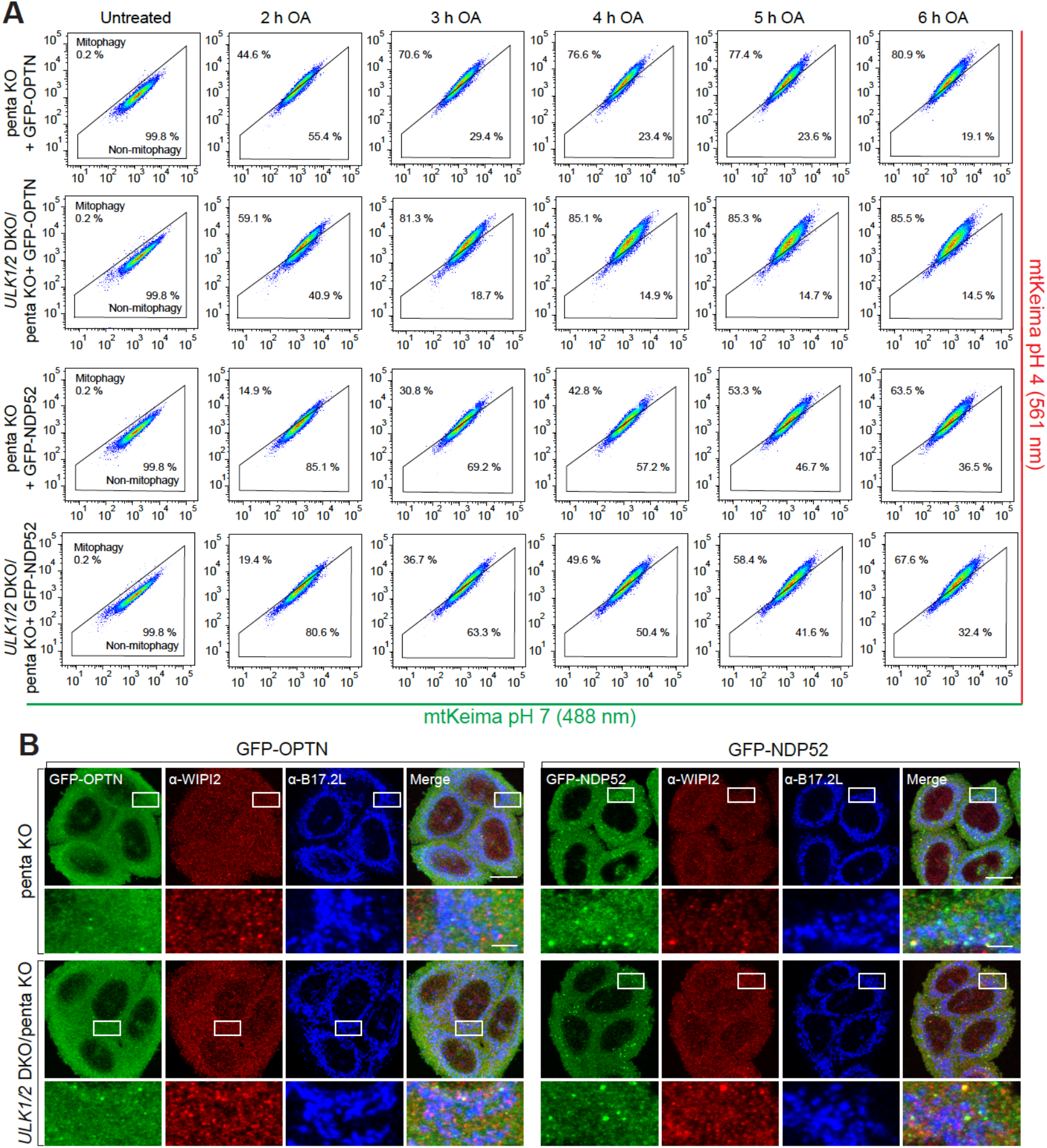
ULK1 and ULK2 are not required for PINK1/Parkin mitophagy. (A) Representative FACS plots for data presented in Figures 1D and 1E. (B) Representative confocal images of untreated penta KO and *ULK1/2* DKO/penta KO expressing BFP-Parkin and GFP-OPTN or GFP-NDP52 immunostained for WIPI2 and B17.2L. Scale bars: overviews, 10 µm; insets, 2 µm.

**Figure S2.**
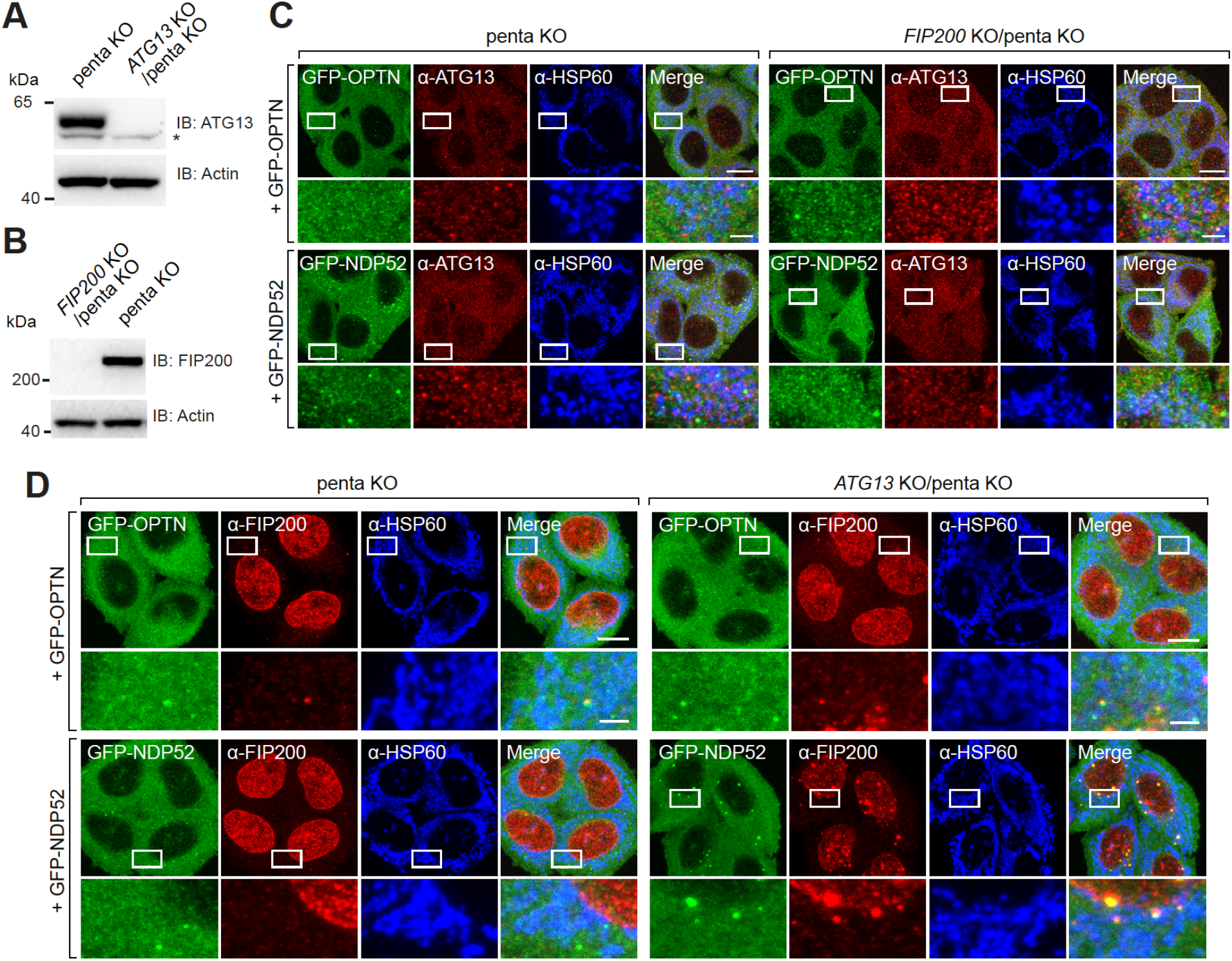
FIP200 and ATG13 are required for PINK1/Parkin mitophagy. (A and B) *ATG13* KO/penta KO (A) and *FIP200* KO/penta KO (B) were confirmed by immunoblotting (IB). kDa: kilodaltons. (C) Representative confocal images of untreated penta KO and *FIP200* KO/penta KO expressing BFP-Parkin and GFP-OPTN or GFP-NDP52 immunostained for ATG13 and mitochondrial HSP60. (D) Representative confocal images of untreated penta KO and *ATG13* KO/penta KO expressing BFP-Parkin and GFP-OPTN or GFP-NDP52 immunostained for FIP200 and mitochondrial HSP60. Scale bars: overviews, 10 µm; insets, 2 µm.

**Figure S3.**
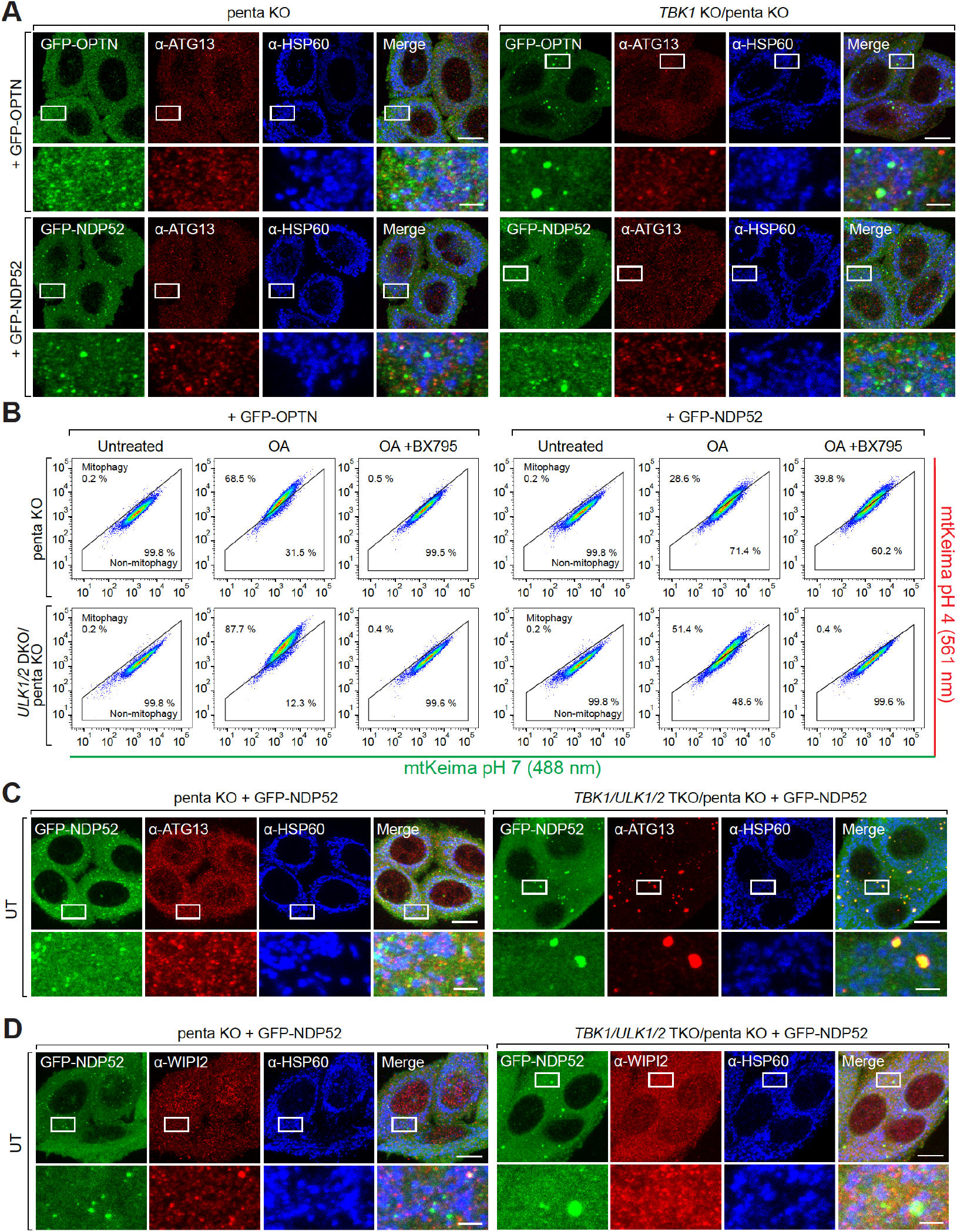
TBK1 is required for OPTN-mediated but not NDP52-mediated mitophagy. (A) Representative confocal images of untreated penta KO and *TBK1* KO/penta KO expressing BFP-Parkin and GFP-OPTN or GFP-NDP52 immunostained for ATG13 and mitochondrial HSP60. (B) Representative FACS plots for data shown in Figures 4A and 4B. (C and D) Representative confocal images of untreated penta KO and *TBK1/ULK1/2* TKO/penta KO expressing BFP-Parkin and GFP-NDP52 immunostained for ATG13 and mitochondrial HSP60 (C) and WIPI2 and mitochondrial HSP60 (D). Scale bars: overviews, 10 µm; insets, 2 µm.

**Figure S4.**
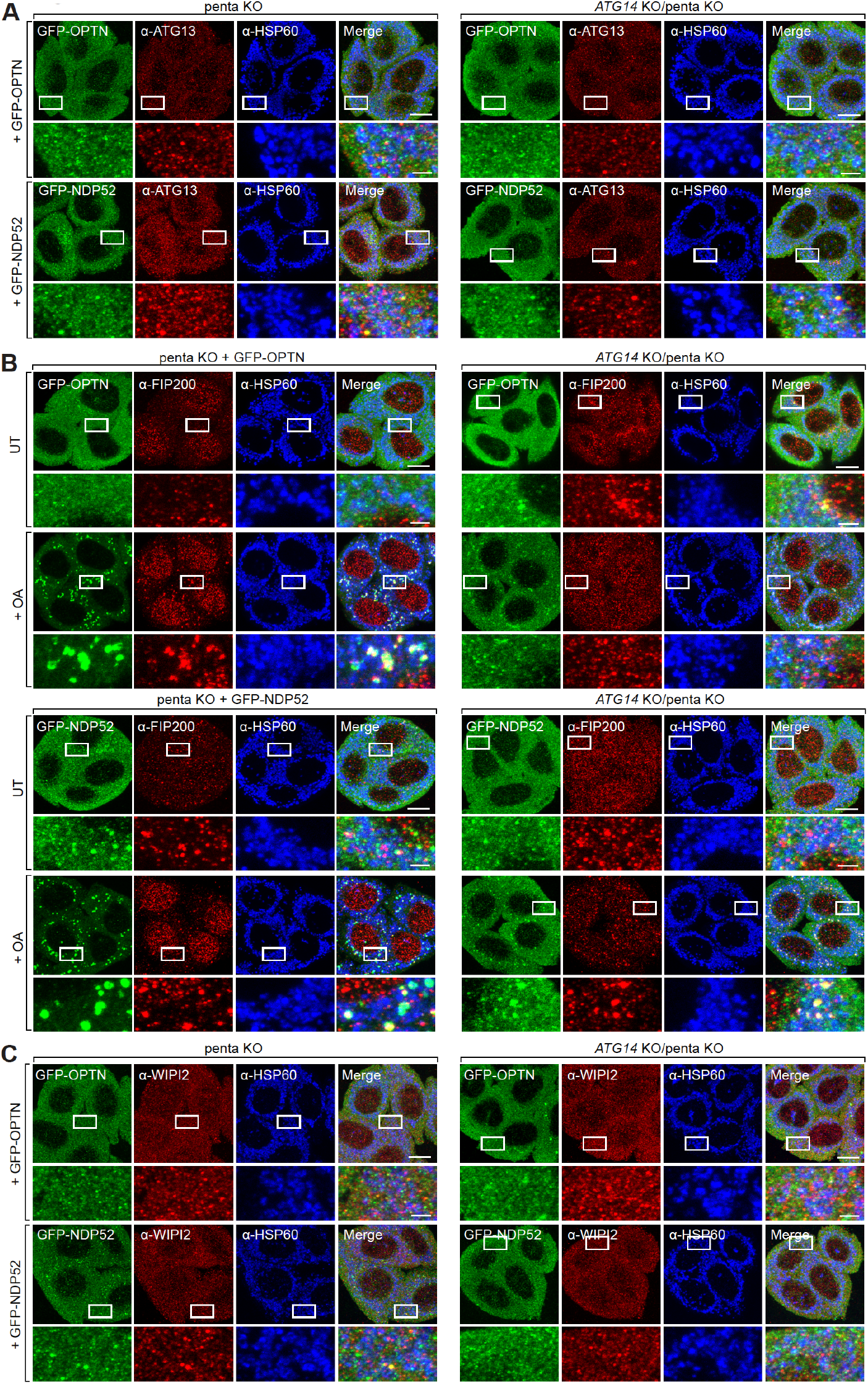
ATG14 is required for PINK1/parkin mitophagy and stable mitochondrial recruitment of ATG13. penta KO and *ATG14* KO/penta KO expressing BFP-Parkin and GFP-OPTN or GFP-NDP52 were left untreated or treated with OA for 1 h as indicated and immunostained for ATG13 and mitochondrial HSP60 (A), FIP200 and mitochondrial HSP60 (B) and WIPI2 and mitochondrial HSP60 (C). Scale bars: overviews, 10 µm; insets, 2 µm.

**Figure S5.**
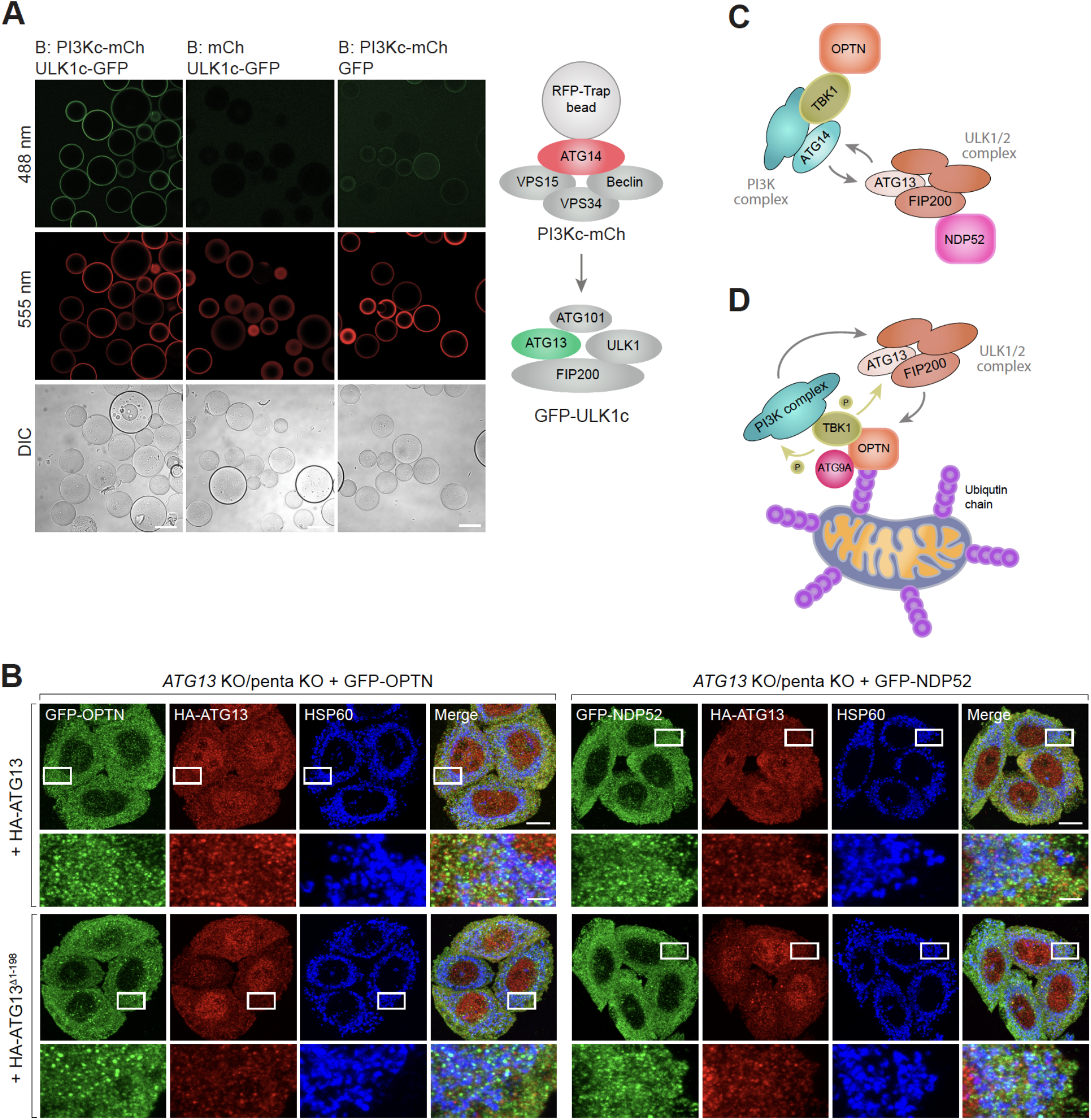
ULK1 complex directly interacts with PI3K complex via HORMA domain of ATG13. (A) Representative confocal images of the recruitment of ULK1 complex labelled with GFP (ULK1c-GFP) to RFP-Trap beads coated with PI3Kc-mCh. (B) Representative confocal images of untreated *ATG13* KO/penta KO expressing BFP-Parkin, GFP-OPTN or GFP-NDP52, and HA-ATG13 or HA-ATG13^.1HORMA^ immunostained for ATG13 and mitochondrial HSP60. (C) Schematic of the initiation complex bound to primary mitophagy adaptors OPTN and NDP52. (D) A model for mitophagy initiation by OPTN. Phosphorylation of the PI3K and ULK1/2 complex might contribute to the process. Scale bars: (A-B) 100 µm; (C) overviews, 10 µm; insets, 2 µm.

**Table S1:**
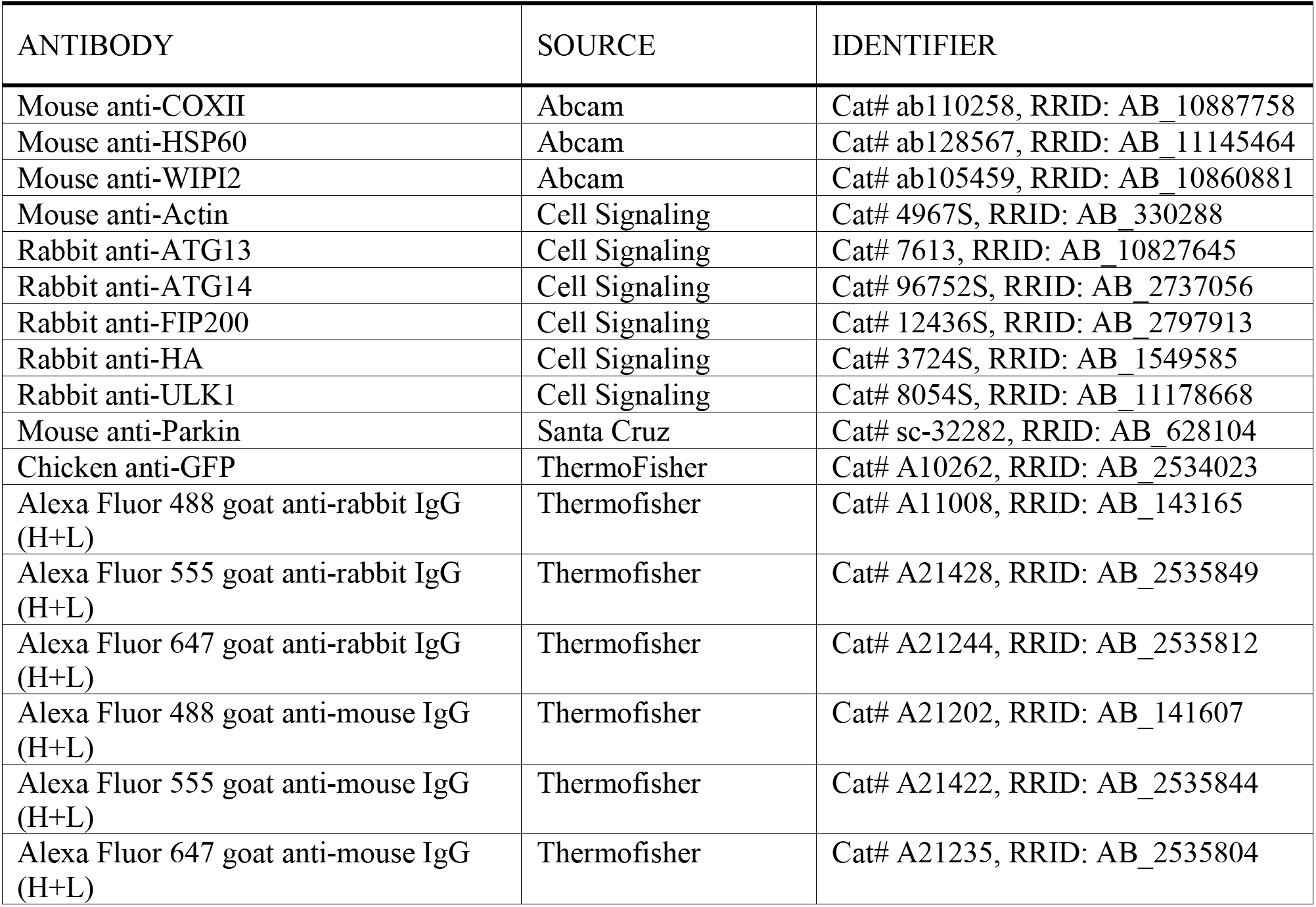
Details of antibodies used in this study.

**Table S2:**
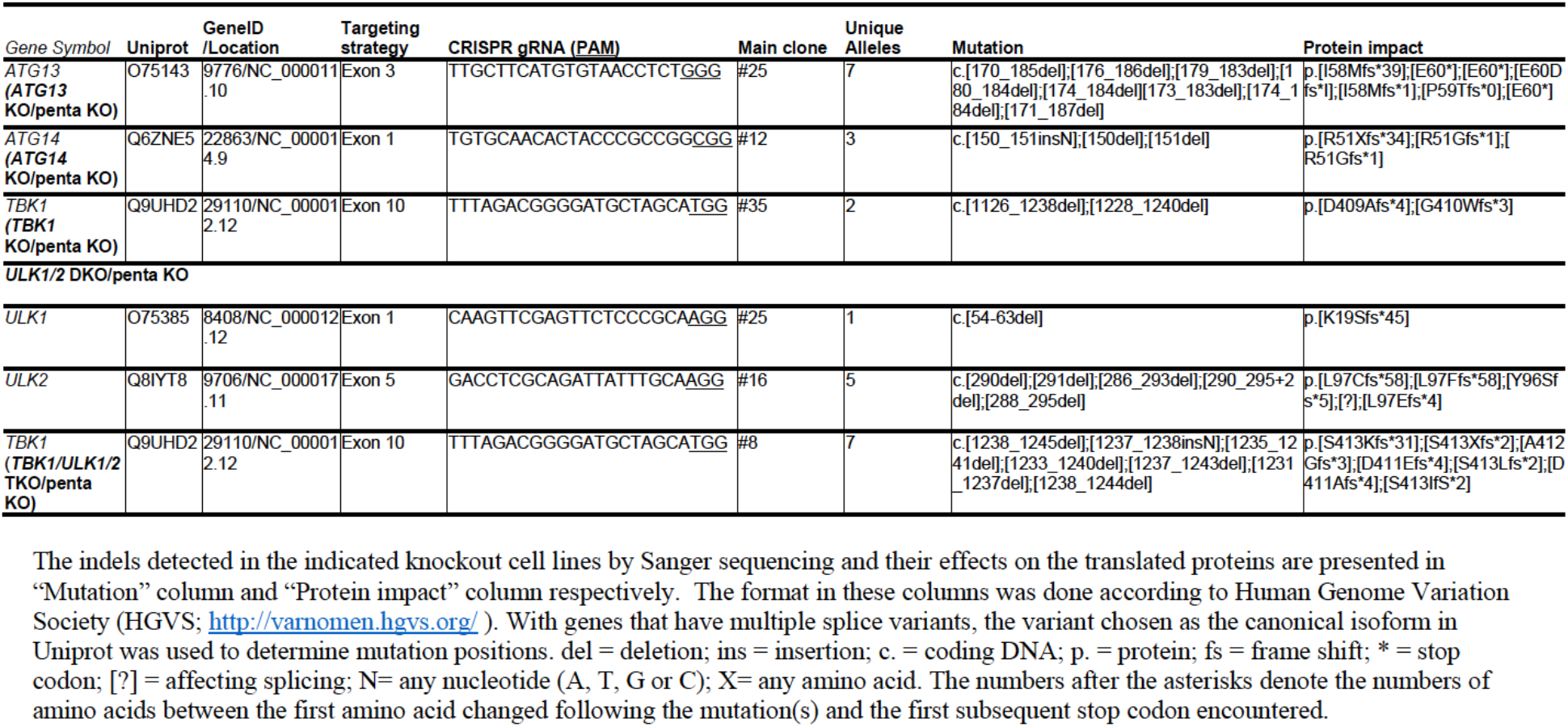
Details of CRISPR sequences, genotyping results of all knockout cell lines in this study.

**Table S3:**
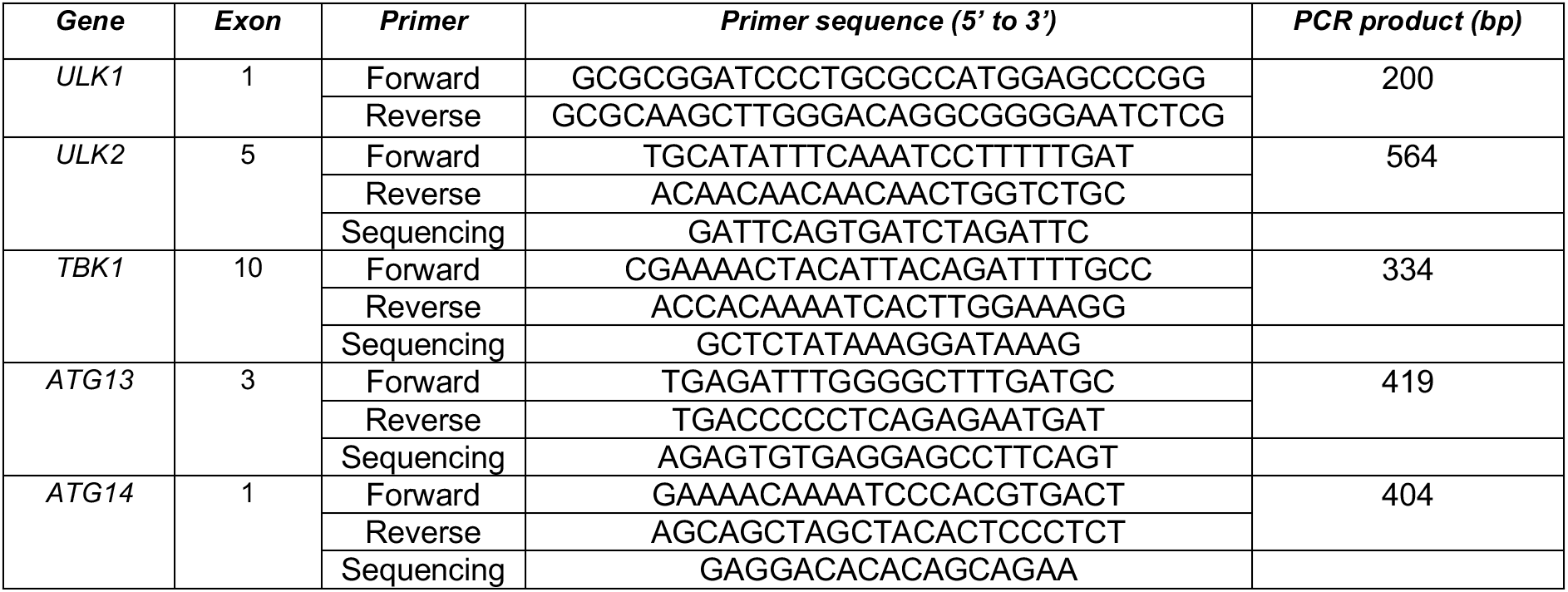
Genotyping primers for sequencing analysis of the generated knockout lines.

**Table S4:**
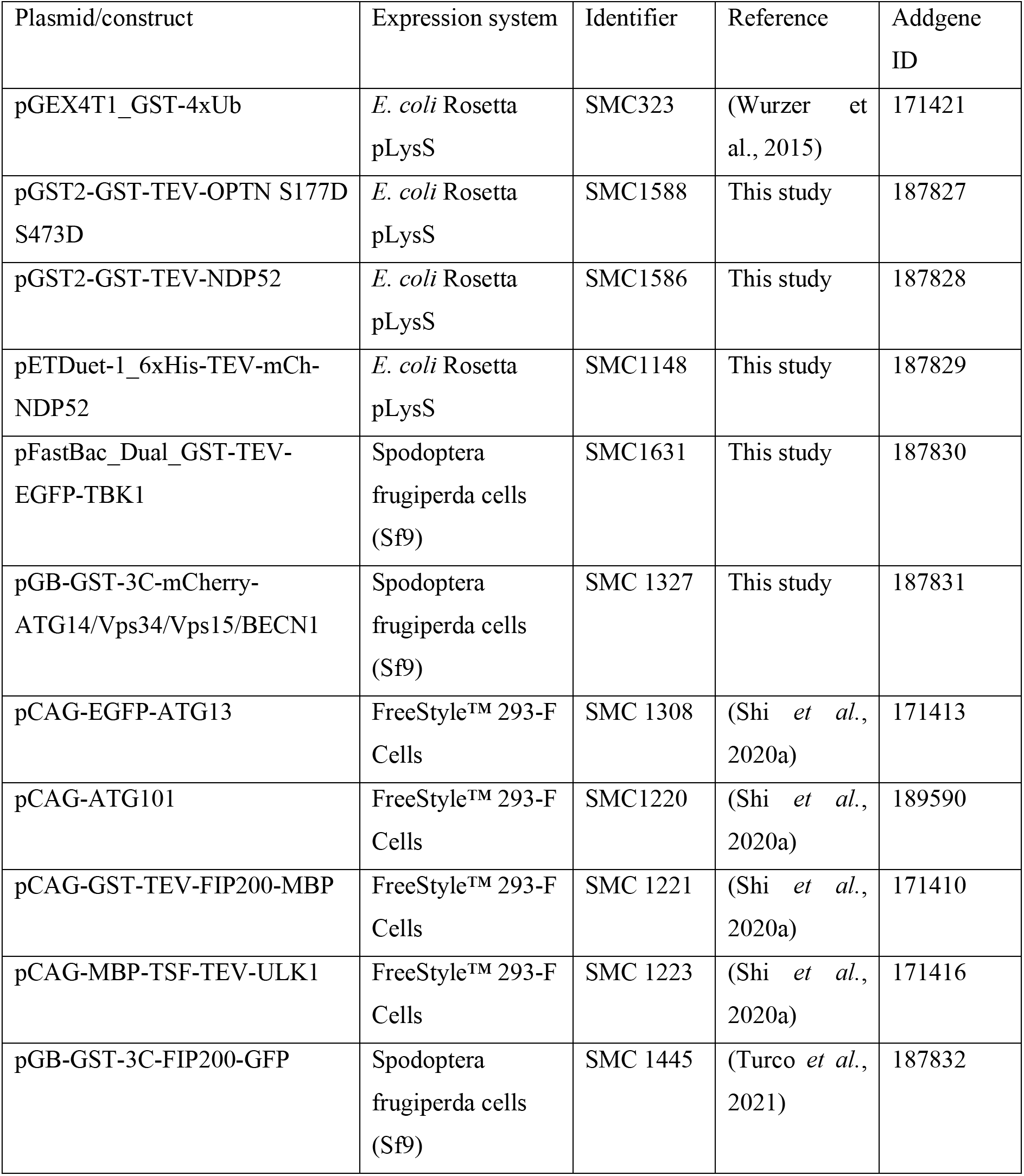
List of expression constructs used in this work.

| Plasmid/construct | Expression system | Identifier | Reference | Addgene ID |
| --- | --- | --- | --- | --- |
| pGEX4T1_GST-4xUb | *E. coli* Rosetta<br>pLysS | SMC323 | (Wurzer et al., 2015) | 171421 |
| pGST2-GST-TEV-OPTN S177D S473D | *E. coli* Rosetta<br>pLysS | SMC1588 | This study | 187827 |
| pGST2-GST-TEV-NDP52 | *E. coli* Rosetta<br>pLysS | SMC1586 | This study | 187828 |
| pETDuet-1_6xHis-TEV-mCh-NDP52 | *E. coli* Rosetta<br>pLysS | SMC1148 | This study | 187829 |
| pFastBac_Dual_GST-TEV-EGFP-TBK1 | Spodoptera<br>frugiperda cells<br>(Sf9) | SMC1631 | This study | 187830 |
| pGB-GST-3C-mCherry-ATG14/Vps34/Vps15/BECN1 | Spodoptera<br>frugiperda cells<br>(Sf9) | SMC 1327 | This study | 187831 |
| pCAG-EGFP-ATG13 | FreeStyle™ 293-F<br>Cells | SMC 1308 | (Shi et al., 2020a) | 171413 |
| pCAG-ATG101 | FreeStyle™ 293-F<br>Cells | SMC1220 | (Shi et al., 2020a) | 189590 |
| pCAG-GST-TEV-FIP200-MBP | FreeStyle™ 293-F<br>Cells | SMC 1221 | (Shi et al., 2020a) | 171410 |
| pCAG-MBP-TSF-TEV-ULK1 | FreeStyle™ 293-F<br>Cells | SMC 1223 | (Shi et al., 2020a) | 171416 |
| pGB-GST-3C-FIP200-GFP | Spodoptera<br>frugiperda cells<br>(Sf9) | SMC 1445 | (Turco et al., 2021) | 187832 |

## Methods

### Antibodies and reagents

The monoclonal and polyclonal antibodies used in this study are listed in Table S1. The polyclonal B17.2L antibody (a kind gift from Prof. Mike Ryan (Monash University)) was previously generated in rabbits using recombinant full length B17.2L antigen (Lazarou et al., 2007).

### Cell culture and generation of knockout lines using CRISPR/Cas9 gene editing

Cell lines in this study (HeLa cell lines and HEK293T) were cultured Dulbecco’s modified eagle medium (DMEM) supplemented with 10 % (v/v) FBS (Cell Sera Australia), 1 % Penicillin-Streptomycin, 25 mM HEPES, 1x GlutaMAX (Life Technologies) and 1x non-essential amino acids (Life Technologies).

Penta KO cells were described previously (Lazarou *et al*., 2015). All knockout cell lines in this study are CRISPR/Cas9-edited cell lines. CRISPR guide RNAs (gRNAs) that target a common exon of all splicing variants of each gene of interest were incorporated into oligonucleotides (Sigma) and introduced into BbsI-linearised pSpCas9(BB)-2A-GFP vector (a gift from Feng Zhang; Addgene # 48138) using Gibson Cloning kit (New England Biolabs). The protocol is available here dx.doi.org/10.17504/protocols.io.j8nlkkzo6l5r/v1. gRNA constructs were then sequence-verified and transfected into penta KO cells with X-tremeGENE 9 (Roche) overnight and single cells that were positive for GFP were sorted by fluorescence activated cell sorting (FACS) into 96 well plates. Single cell colonies were then expanded and collected for screening by immunoblotting for the loss of the targeted gene product. Once the knockout cell lines were identified, gene editing was detected using Sanger sequencing. The CRISPR-targeted regions were amplified via PCR and PCR products were sequenced using sequencing primers that anneal to the amplified regions (Table S3). For analysis purpose, the same regions in the parental line were also sequenced. Synthego ICE v2 CRISPR Analysis Tool (synthego.com/products/bioinformatics/crispr-analysis) was used to compare the sequencing data from the control and the knockout cells, allowing the identification of gene edits in the knockout cells (Table S2). For *ULK1* KO cell lines (in *ULK1/2* DKO/penta KO and *TBK1/ULK1/2* TKO/penta; Table S2), sequencing the amplified PCR products appeared to be difficult, we therefore cloned them into a pGEM4Z vector prior to sequencing analysis (see Table S3 for details of genotyping primers). Although the sequencing reads were not of high quality in some parts, there was a clear edit of 10 bp deletion at the CRISPR target region which resulted in frameshift and premature stop codon (Table S2). Nevertheless, there was no protein detected by immunoblotting (Figure 1A). Where antibodies were not available, putative KO clones were first identified by three primer PCR (Yu et al., 2014) and the presence of frameshift indels was determined by Sanger sequencing. The protocol is available online dx.doi.org/10.17504/protocols.io.8epv59yx5g1b/v1. Multiple knockout lines were generated by sequential transfections of one or multiple gRNA constructs. For *ULK1/2* DKO/penta KO, ULK1 CRISPR was transfected into penta KO cells to make *ULK1* KO/penta KO. The introduction of ULK2 CRISPR into *ULK1* KO/penta KO produced *ULK1/2* DKO/penta KO. Subsequent editing of *ULK1/2* DKO/penta KO with TBK1 CRISPR generated *TBK1/ULK1/2* TKO/penta KO.

### Cloning and generation of stable cell lines

pBMN-mEGFP-OPTN (Addgene #188784), pBMN-mEGFP-NDP52 (Addgene # 188785), pBMN-BFP-Parkin (Addgene #186221) and universal vector backbones pBMN-mEGFP (Addgene #188643), pBMN-HA-C1 (Addgene #188645) and pBMN-BFP-C1 (Addgene #188644) was previously described (Nguyen *et al*., 2021; Padman *et al*., 2019). Open reading frames (ORFs) of ATG13 and ATG13^.11-198^ referred to as ATG13^.1HORMA^ were introduced into linearised pBMN-HA-C1 using Gibson Cloning kit (New England Biolabs) to create pBMN-HA-ATG13 and ATG13^.1HORMA^ (Addgene #186223 and #18622 respectively). All constructs were sequence-verified.

Stable cell lines were generated using retroviral system as described previously (Nguyen *et al*., 2021). First, a retroviral vector (pCHAC-mito-Keima and all pBMN constructs) containing an ORF of interest and retroviral packaging plasmids (VSV-G and Gag-Pol) were transfected into HEK293T cells using Lipofectamine LTX (Life Technologies) for 15 h. The next day, the transfection media was replaced with fresh growing DMEM to allow the assembly and secretion of retroviruses. After 24 h incubation, supernatant containing retroviruses was collected, filtered and applied onto the desired HeLa cells for 24-48 h in the presence 8 mg/ml polybrene (Sigma). Cells were then recovered and expanded in growing DMEM for 5-7 days prior to sorting by FACS to match protein expression levels where possible.

### Mitophagy treatment and Immunoblotting

Translocation and mitophagy experiments were achieved by incubating cells with 10 μM Oligomycin (Calbiochem), 4 μM Antimycin A (Sigma) and 10 μM QVD (ApexBio) in full growth medium for different time points as indicated in figure legends. For the analysis of BX795 treatment, the indicated samples were pre-treated for 30 min with 1 µM BX795 in growth media before inducing mitophagy in the presence of 1µM BX795.

For Western blot analysis, cells from 6 well plates were washed with 1x PBS, harvested using cell scrapers and lysed in 1× LDS sample buffer (Life Technologies) supplemented with 100 mM dithiothreitol (DTT; Sigma). Samples were heated at 99 °C with shaking for 7-10 min. Approximately 25-80 μg of protein per sample was analysed by 4-12% Bis-Tris gels (Life Technologies) according to manufacturer’s instructions. Gels were electro-transferred to polyvinyl difluoride membranes (PVDF) and immunoblotted using indicated antibodies. The protocol is deposited here dx.doi.org/10.17504/protocols.io.kxygxzxr4v8j/v1

### Immunofluorescence assay and confocal image analysis

For immunofluorescence, 48 h before treatment cells were seeded on HistoGrip (ThermoFisher) coated glass coverslips for. All following steps were performed at room temperature. Samples were first fixed with 4% (w/v) paraformaldehyde (PFA) in 0.1 M phosphate buffer on a rocker for 10 min. Samples were then rinsed three times with 1x PBS, permeabilized with 0.1% (v/v) Triton X-100 in 1x PBS (10 min) and then blocked with 3% (v/v) goat serum in 0.1% (v/v) Triton X-100/1x PBS (15 min). The samples were incubated with indicated primary antibodies made up in 3 % (v/v) goat serum in 0.1 % (v/v) Triton X-100/1x PBS for 90 min and rinsed three times with 1x PBS. Following 1 h incubation with secondary antibodies conjugated to AlexaFluor-488, Alexa-Fluor-555, AlexaFluor-633, or AlexaFluor-647 (ThermoFisher), the coverslips were washed three times with 1x PBS and mounted with a TRIS buffered DABCO-glycerol mounting medium onto glass slides. Imaging of the coverslips wwas done with an inverted Leica SP8 confocal laser scanning microscope under 63x/1.40NA objective (Oil immersion, HC PLAPO, CS2; Leica microsystems). Images were acquired in 3D by optical sectioning using an with a minimum z-stack range of 1.8 µm and a maximum voxel size of 90 nm laterally (x,y) and 300 nm axially (z) using a Leica HyD Hybrid Detector (Leica Microsystems) and the Leica Application Suite X (LASX v2.0.1). All images are displayed as z-stack maximum projections. For image analysis, three images were taken per sample at random stage positions. If the field-of -view of any image contained less than 30 cells, another random one was chosen for imaging. The protocol is described here dx.doi.org/10.17504/protocols.io.5qpvobz99l4o/v1

Automated 3D image segmentation approach with 3D ROI manager (v3.93) and FeatureJ (v3.0.0) plugins for FIJI (v1.52p) was employed to process and analyse all 3D image data. As described previously (Padman *et al*., 2019), the image analysis process includes three steps: object detection, object measurement and analysis. To account for the lack of foci in the untreated conditions, ATG13 and WIPI2b foci were detected and segmented using a global thresholding strategy. For each experiment repeat, a montage of maximum intensity projections of all images was assembled, and new minimum and maximum intensity values required for linear histogram normalization of that montage were calculated. The minimum and maximum intensity values were defined if the number of pixels at a given intensity exceeded 0.02 % of all pixels in the montage. The histogram of each individual image was then rescaled between these new maximum and minimum values to generate normalized data. This data was then subjected to 3D noise filtering (3D median filter; 1.5 voxels) and extraction of 3D ROIs using a global threshold (ATG13 and WIPI2b, 64; arbitrary) via simple 3D thresholding (300 voxel maximum size). Re-applying each extracted 3D ROI to the original corresponding image allowed the determination of volumes and fluorescence intensities for each segmented object in all available channels. All the measurements and ROIs were saved for subsequent analyses. Foci volume and number for ATG13 and WIPI2b were calculated by direct measurement of these parameters from the 3D ROIs. The number of cells in each image was counted manually.

### Mito-Keima autophagosome-lysosome fusion mitophagy assay

Cells were seeded into 24 well plates one day before the experimental day. Following mitophagy induction at indicated time points, cells were washed with 1x PBS, detached with trypsin (Life Technologies) and harvested in normal growth media. Samples were then centrifuged at 4 °C and resuspended in ice cold sorting buffer containing 10 % v/v FBS and 0.5 mM EDTA in 1x PBS). Samples were analysed using the FAVSDiva software on a LSR Fortessa X-20 cell sorter (BD Biosciences). Lysosomal mtKeima was measured using dual excitation ratiometric pH measurements at 488 (pH 7) and 561 (pH 4) nm lasers with 695 nm and 670 nm detector filters respectively. Additional channel used was GFP (Ex/Em; 488 nm/530 nm) for fluorescence compensation. For each sample, 30,000 events were collected, and data were analysed using FlowJo (version 10). The protocol is deposited online dx.doi.org/10.17504/protocols.io.q26g74e1qgwz/v1.

### Transmission Electron Microscopy (TEM) imaging

Samples were fixed with pre-warmed 4 % PFA in 0.1 M phosphate buffer (pH 7.2) at 37 °C for 1 hour followed by overnight post-fixation with 2.5 % glutaraldehyde in 0.1 M sodium cacodylate buffer (pH 7.4) at 4 °C. Samples were rinsed three times with 0.1 M sodium cacodylate buffer. All subsequent sample preparation steps were microwave assisted using a BioWave Pro microwave system (Pelco). The samples were osmicated with 1 % (w/v) OsO4, 1.5 % (w/v) K3Fe(CN)6 in 0.1 M cacodylate buffer for 1 hour at 4 °C, before exposure to three microwave duty-cycles (120 s on, 120 s off) at 100 W under vacuum. The samples were rinsed three times with MilliQ water and en bloc stained with 2 % (w/v) aqueous uranyl acetate with three 100 W microwave duty-cycles (120 s on, 120 s off) under vacuum. Samples were then rinsed with MilliQ water twice before the cells were scraped, resuspended in 500 µL MilliQ, and pelleted (10,000x g). The cells were then repelleted in 500 µL 70% ethanol, and further resuspended and pelleted in 300 µL low-melting point agarose at 37°C. The solidified agarose-cell pellet was divided into 1 mm cubes using a razor blade for subsequent processing. Microwave assisted dehydration was performed by graduated ethanol series (80 %, 90 %, 95 %, 100 %, 100 % (v/v); each at 150 W for 40 s) and propylene oxide (100 %, 100 % (v/v); each at 150 W for 40 s). The samples were infiltrated with Araldite 502/Embed 812 by graduated concentration series in propylene oxide (25 %, 50 %, 75 % 100 %, 100 % (v/v); 180 s at 250 W under vacuum), then polymerized at 60 °C for two days. Ultracut UCT ultramicrotome (Leica Biosystems) equipped with a 45 °C diamond knife (Diatome) was used to section the resin embedded samples, and 75 nm ultrathin sections were collected and loaded on 75 mesh copper grids. The grids were stained at room temperature using 2 % (w/v) aqueous uranyl acetate (5 min) and Reynolds lead citrate (3 min) before routine imaging on a JEM-1400PLUS TEM (JEOL). For quantification, two ultrathin sections per sample were surveyed and 15 cells were randomly selected at a low magnification (2,000x) for further analysis. Cells were excluded from selection if they were directly adjacent to a previously selected cell, partially obstructed by a grid bar, or lacking a visible nucleus. Valid cells were imaged by acquiring a montage of high magnification (5,000X) images, to enable the cataloguing of all visible autophagosomal structures within the cell. All the visible autophagosomal structures were further imaged at 10,000X magnification for validation. The protocol is deposited here https://dx.doi.org/10.17504/protocols.io.14egn75zmv5d/v1

### Protein expressions and purification

Linear tetra-ubiquitin (GST-4xUb) was expressed and purified as previously described (Wurzer *et al*., 2015). The protocol is deposited online (dx.doi.org/10.17504/protocols.io.bvjbn4in)

**GFP-labelled ULK1 complex was expressed in** FreeStyle™ 293-F Cells by transient transfection. The cells grown at 37°C in FreeStyle™ 293 Expression Medium (Thermo, 12338-026) were seeded to density of 0.7 x 10^6 cells per ml the day before transfection. On the day of transfection, 400 ml culture at density of 1×10^6 cells per ml was transfected by addition of vortexed transfection mixture. The mixture consisted of two components: 400 ug of the MAXI-prep DNA that was pre-diluted in 13 ml of Opti-MEM® I Reduced Serum Medium (Thermo, 31985-062) and 800 ug Polyethylenimine (PEI 25K, Polysciences CatNo 23966-1) likewise pre-diluted in 13 ml of Opti-MEM media. 24 hours post transfection, the culture was fed by addition of 100 ml of EX-CELL® 293 Serum-Free Medium (SigmaAldrich, 14571C-1000ML). 48H post transfection, the cells were harvested by centrifugation at 270 g for 20 minutes, washed by 1xPBS and flash-frozen in liquid nitrogen prior to storage at −80°C. The ULK1 complex was previously purified (Shi *et al*., 2020a). The protocol is available online (dx.doi.org/10.17504/protocols.io.bvn2n5ge).

FIP200-GFP was expressed in Spodoptera frugiperda cells (Sf9) and purified as described previously (Turco *et al*., 2021). The protocol is available online (dx.doi.org/10.17504/protocols.io.dm6gpbkq5lzp/v1).

The pGST2-GST-TEV-OPTN(S177D S473D) and pGST2-GST-TEV-NDP52 constructs were generated by introducing TEV cleavage site to corresponding vectors (Chang *et al*., 2021) (Protocol: dx.doi.org/10.17504/protocols.io.bp2l61znrvqe/v1). The proteins were expressed in *E. coli* Rosetta pLySS cells. Cells were grown in LB medium at 37°C until an OD600 of 0.4. Next, the culture was brought to 18°C and grown to an OD600 of 0.8. Protein expression was induced with 100 μM IPTG and grown for further 16 h at 18°C. Cells were pelleted and resuspended in a buffer containing 50 mM HEPES, pH 7.5, 300 mM NaCl, 2 mM MgCl2, 2 mM β-mercaptoethanol, cOmplete protease inhibitors (Roche), and DNase. Cells were lysed by freeze thawing and 2 × 30-s sonication. Lysates were cleared by ultracentrifugation (25,000 rpm for 30 min at 4°C in a Ti45 rotor). Supernatant was incubated with 5 ml Glutathione Sepharose 4B beads slurry (Cytiva) for 1h at 4°C. Beads were then washed with low salt (50 mM HEPES, pH 7.5, 300 mM NaCl, and 1 mM DTT) buffer, followed by high salt (50 mM HEPES, pH 7.5, 500 mM NaCl, and 1 mM DTT) and low salt buffers. Finally, beads were incubated overnight with TEV protease at 4°C. The supernatant containing cleaved protein was filtered through a 0.45 μm syringe filter, concentrated down to 0.5 ml and applied onto a Superdex 200 column (10/300 Cytiva) pre-equillibrated with a buffer containing 25 mM HEPES, pH 7.5, 150 mM NaCl, and 1 mM DTT. Fractions containing pure proteins were pooled, concentrated, snap frozen in liquid nitrogen, and stored at −80°C.

The mCherry-NDP52 gene coding sequence was cloned in a pET-Duet1 vector, with a N-terminal 6xHis tag followed by a TEV cleavage site. The protein was expressed in *E. coli* Rosetta pLySS cells. Cells were grown in LB medium at 37°C until an OD600 of 0.4. Next, the culture was brought to 18°C and grown to an OD600 of 0.8. Protein expression was induced with 50 μM IPTG and grown for a further 16 h at 18°C. Cells were pelleted and resuspended in a buffer containing 50mM HEPES pH7.5, 300 mM NaCl, 1 mM MgCl2, 10 mM Imidazole, 2 mM Beta-Mercaptoethanol, cOmplete protease inhibitors (Roche), and DNase (Sigma). Cells were lysed by freeze thawing and 3 × 30 s sonication. Lysates were cleared by ultracentrifugation (40,000 rpm for 45 min at 4°C in a Ti45 rotor). Supernatant was filtered (0.45 μm) applied to a 5-ml His-Trap HP column and eluted via a stepwise imidazole gradient (50, 75, 100, 150, 200, and 300 mM). Fractions at 75–100 mM imidazole containing His-TEV-mCh-NDP52 were pooled and subjected to TEV protease cleavage over night at 4°C. The cleaved protein was concentrated using a 50kDa cut-off Amicon filter and injected to a Superdex200 16/600 column (Cytiva) and eluted with a buffer containing 25mM HEPES pH7.5, 150mM NaCl, 1mM DTT. Purified protein was concentrated, snap frozen in liquid nitrogen, and stored at −80°C. The protocol is deposited online (dx.doi.org/10.17504/protocols.io.5qpvobdr9l4o/v1)

To generate GFP-TBK1 constructs the insect codon optimized TBK1 gene was purchased from GenScript and cloned with respective tags into pFastBac_Dual (Table S4). Generated constructs were used for expression in Sf9 insect cells using the Bac-to-Bac system. The bacmid DNAs (2.5 μg per construct), obtained by amplification in DH10BacY cells were used to transfect Sf9 insect cells using FuGene transfection reagent (Promega). About 7 days after transfection the V0 virus was harvested and used to produce a V1 virus stock, which in turn was used to infect 1 L Sf9 cells (1 million/ml) for protein expression. After infection cells were monitored and harvested by centrifugation when they reached a viability of 95–98%. Cell pellets were washed with PBS, flashfrozen in liquid nitrogen, and stored at −80 °C until purification. For purification of GFP-TBK1, a cell pellet corresponding to 1L culture was thawed and resuspended in 30 ml lysis buffer (50 mM Tris-HCl pH 7.4, 300mM NaCl, 2 mM MgCl2, 5% glycerol, 2 mM β-Met, 1 μl Benzonase (Sigma), CIP protease inhibitor (Sigma), cOmplete EDTA-free protease inhibitor cocktail (Roche)). Cells were additionally disrupted with Dounce homogenizer. Lysates were cleared by centrifugation (19 000 rpm for 45 min at 4°C in a Fiberlite F21-8×50y (Thermo Scientific). Supernatant was incubated with 5 ml Glutathione Sepharose 4B beads slurry (Cytiva) for 2h at 4°C. Then beads were washed five times with 50 mM Tris-HCl pH 7.4, 300 mM NaCl, 5% glycerol, 1 mM DTT and incubated overnight with TEV protease at 4°C. The supernatant containing cleaved protein was filtered through a 0.45 μm syringe filter, concentrated down to 0.5 ml and applied onto a Superdex 200 column (10/300, Cytiva) pre-equillibrated with a buffer containing 20 mM Tris-HCl pH 7.4, 300 mM NaCl, and 1 mM DTT. Fractions containing pure proteins were pooled, concentrated, snap frozen in liquid nitrogen, and stored at −80°C. The protocol is available at dx.doi.org/10.17504/protocols.io.81wgb6wy1lpk/v1

Genes coding for protein sequences of human ATG14, BECN1, VPS34, VPS15 were codon optimized for the Sf9 insect cell expression system, and synthetic genes were purchased from GenScript. All four ORFs were insertent to a GoldenBac plasmid (pGB) via Golden Gate approach by the Vienna BioCenter Core Facilities (VBCF) Protech Facility. To express mCherry labelled PI3KC3-c1 1 L culture of Sf9 cells growing in Sf921 medium at 1-1.5 mil/ml cells/volume were infected with 1 ml of Virus 1 (V1). Baculovirus was obtained by transfection of Sf9 cells with a policystronic construct coding for the mCherry labelled PI3KC3-C1 complex (Table S4). After infection cells were monitored and harvested by centrifugation when they reached a viability of 95-98%. Cell pellets were washed with PBS, flash frozen in liquid nitrogen, and stored at −80 °C until purification. To purify mCherry labelled PI3KC3-c1 the cell pellet corresponding to 1L culture was thawed and resuspended in 50 ml lysis buffer (50 mM HEPES pH 7.5, 300 mM NaCl, 0.5 % CHAPS, Benzonase, 1 mM MgCl_2_, 1 mM DTT, CIP protease inhibitor (Sigma), cOmplete EDTA-free protease inhibitor cocktail (Roche). Cells were additionally disrupted with Dounce homogenizer. Lysates were cleared by centrifugation at 25 000 rpm for 45 min at 4°C in a Ti45 rotor (Beckman). Supernatant was incubated with 5 ml of Glutathione Sepharose 4B beads slurry (Cytiva) for 1h at 4°C. Then beads were washed twice with 50 mM HEPES pH 7.5, 30 0mM NaCl, 0.5% CHAPS, 1 mM DTT, twice with 50 mM HEPES pH 7.5, 700 mM NaCl, 1 mM DTT and finally twice with 50 mM HEPES pH 7.5, 300 mM NaCl, 1 mM DTT. To elute the protein complex the beads were incubated overnight with Precission (3C) protease at 4°C. The supernatant containing cleaved protein was filtered through a 0.2 μm syringe filter, concentrated down to 0.5 ml and applied onto a Superose 6 column (10/300, Cytiva) pre-equillibrated with a buffer containing 50 mM HEPES pH 7.5, 200 mM NaCl, 1 mM DTT. Fractions containing pure proteins were pooled, concentrated, snap frozen in liquid nitrogen, and stored at −80°C. The protocol is available at dx.doi.org/10.17504/protocols.io.8epv59mz4g1b/v1.

### Microscopy-based bead protein-protein interaction assay

Glutathione Sepharose 4B beads (GE Healthcare), RFP- or GFP-Trap beads (ChromoTek), were mixed with GST, mCherry or GFP tagged bait proteins, respectively to the final concentrations of 10 μM (GST-4xUb), 2.5 μM (GFP-TBK1) and 1 μM (PI3Kc1-mCh). Beads were incubated with bait proteins at 4 °C for 1 h, and then washed twice with washing buffer (25mM TRIS-Cl pH 7.5, 150mM NaCl, 1mM DTT). 1 μl of respective beads was transferred into the well of a 384-well glass-bottom microplate (Greiner Bio-One) pre-filled with 20 μl of 25mM TRIS-Cl pH 7.5, 150mM NaCl, 1mM DTT and prey proteins. For the experiments shown in Figure 2G, mCh-NDP52 and FIP200-GFP were used at the final concentration of 100 nM and 500 nM, respectively. For the experiments shown in Figures 6A and 6B, PI3Kc1-mCh and mCherry were used at the final concentration of 200 nM, GFP-TBK1 and OPTN S177D S473D at the final concentration of 500 nM. For the experiments shown in Figures 6C, S5A and S5B, ULK1c-GFP, GFP and NDP52 were used at the final concentration of 500 nM, and PI3Kc1-mCh at the final concentration of 200 nM. The samples were incubated for 30 min and imaged at the equilibrium by Zeiss LSM 700 confocal microscope equipped with Plan Apochromat 20X/0.8 WD 0.55 mm objective.

For quantification of mCh-NDP52 recruitment to the beads in the presence and absence of FIP200-GFP (Figure 2I) using ImageJ software eight lines were drawn across each bead and the maximum brightness value along each the line was taken. Next, the average brightness of an empty area of each picture was measured (background fluorescence) and subtracted from the maximal fluorescence for each bead. The average values for each sample were averaged between 3 independent replicates and plotted with the relative standard errors. For the quantification described above, statistical analysis was performed. Statistical significance of the difference between 2 samples was established by 2 samples unpaired t test. Significant differences are indicated with * when p value ≤ 0.05, ** when p value ≤ 0.01, *** when p value ≤ 0.001. The Protocol for this is available online (dx.doi.org/10.17504/protocols.io.6qpvr6nwpvmk/v1)

## Acknowledgments

We would like to thank the laboratory of Noboru Mizushima (The University of Tokyo) for sharing *FIP200* KO/penta KO cells and the laboratory of Mike Ryan (Monash Biomedicine Discovery Institute, Monash University) for sharing anti-B17.2L antibodies. We also thank Monash Flow Cytometry Platform (FlowCore), the Monash Micro Imaging Platform, the Ramaciotti Centre for Cryo-Electron Microscopy and the Max Perutz Labs BioOptics facility for technical support. This work was supported by the National Health and Medical Research Council (NHMRC) (GNT1106471 to M.L.), the Australian Research Council (ARC) Discovery Project (DP200100347 to M.L.), the combined efforts of the Michael J Fox Foundation for Parkinson’s Research (MJFF) and Aligning Science Across Parkinson’s (ASAP) initiative (ASAP-000350 to M.L. and S.M.), the Human Frontiers Science Program RGP0026/2017 (to S.M.) and a research and travel grant from Flanders Fund for Scientific Research (FWO-Flanders, Project number: 1228021N and V40192N to E.A.).

## Author contributions

T.N.N. and M.L. conceived the projects; T.N.N, J.S.M, S.M. and M.L. designed experiments; T.N.N, J.S.M, G.K., W.K.L, E.A., D.F., S.S., B.S.P., M.S., R.S.J.L. performed experiments. T.N.N. and M.L. wrote the manuscript and all authors contributed to preparing and editing the manuscript.

## Data availability

All data are provided within the manuscript and supplement.

